# The ID93 + GLA-3M-052-LS vaccine candidate elicits mucosal and systemic immunogenicity and protective efficacy against *Mycobacterium tuberculosis* challenge in BCG-primed Collaborative Cross inbred mice

**DOI:** 10.1101/2025.10.31.685426

**Authors:** Emily A. Voigt, Anas Alsharaydeh, Darshan N. Kasal, Madeleine F. Jennewein, Devin S. Brandt, Susan Lin, Jasneet Singh, Julie Bakken, Raodoh Mohamath, Pauline Fusco, Jordi B. Torrelles, Gillian Beamer, Christopher B. Fox

## Abstract

New vaccine approaches are needed against tuberculosis (TB). We sought to optimize mucosal immunogenicity and protective efficacy by modulating the adjuvant component and route of immunization of a next-generation TB vaccine using the recombinant TB vaccine antigen (Ag) ID93. ID93-specific mucosal and systemic immunogenicity and protective efficacy were assessed in the Collaborative Cross 004 mouse strain, a mouse strain susceptible to *Mycobacterium tuberculosis* (*Mtb*) infection, as a suitable model of *Mtb* susceptible populations. Immunogenicity data from various vaccine candidates were used to select lead vaccine candidates with the most preferred immunostimulatory profiles using a pre-determined desirability index. A liposomal adjuvant system containing synthetic TLR4 and TLR7/8 ligands (GLA-3M-052-LS), administered by a heterologous intramuscular-intranasal regimen, induced an optimal comprehensive immune response profile including high levels of mucosal antibody and Th1 CD4^+^ T cells in the lungs. In BCG-primed mice, immunization with intramuscular followed by intranasal ID93 + GLA-3M-052-LS boosts significantly reduced *Mtb* burden in the lungs after challenge *vs.* BCG vaccinated mice alone. Thus, ID93 + GLA-3M-052-LS represents a promising next-generation TB vaccine candidate suitable for testing in additional preclinical models.

## INTRODUCTION

The only approved vaccine against the development of tuberculosis (TB) disease, *Mycobacterium bovis* Bacille Calmette *et Guérin* (BCG), is widely used in children; however, efficacy in adolescents and adults is low, and BCG-vaccinated TB patients can still transmit *Mycobacterium tuberculosis* (*Mtb*).^1^ *Mtb* transmission from an active TB case varies according to settings, but the reproductive number is reported to be as high as 4.3.^2^ In addition, the discovery of sub-clinical TB cases without disease symptoms,^3^ ensures a continued prevalence of new *Mtb* infections if new countermeasures in detection and prevention are not developed. For these reasons, TB vaccine formulations and immunization regimens that can generate mucosal immune responses in the lungs, with increased potential to reduce TB disease, infection, and/or transmission are of particular interest.

The leading TB vaccine in clinical development, consisting of the recombinant protein antigen (Ag) M72 with the liposomal adjuvant system AS01E, demonstrated 50% protective efficacy from disease in a Phase 2b trial and a large Phase 3 efficacy trial initiated in 2024.^4^ Another vaccine candidate consisting of a recombinant protein with adjuvant, ID93 + GLA-SE, has demonstrated promising safety and immunogenicity in Phase 1/2a clinical testing and is available as a thermostable lyophilized presentation.^5^

Both of these current vaccine candidates are administered by the intramuscular (i.m.) route and include Toll-like receptor (TLR) 4 agonists; thus, considerable room remains to explore and develop vaccines using other adjuvant types and methods of delivery. To build on the success of these vaccine candidates, adjuvants that target additional pattern recognition receptors and are suitable for mucosal delivery are of particular interest.

We paired ID93 [composed of the four *Mtb* Ags encoded by genes Rv3619, Rv1813, Rv3620, and Rv2608 linked in tandem] with several new adjuvant candidates and alternative immunization route regimens to generate enhanced mucosal and systemic immune responses in Collaborative Cross (CC) 004 mice. As a benchmark, the novel vaccine formulations were compared to ID93 + GLA-SE. CC004 mice were selected for their greater susceptibility to *Mtb* infection, a consequence of BCG providing lower protective efficacy compared to C57BL/6 mice,^6^ therein representing a potentially more suitable model of *Mtb*-susceptible populations most in need of vaccine-mediated protection.

Our results indicate that ID93 + GLA-3M-052-LS adjuvant administered by a heterologous i.m. and intranasal (i.n.) route regimen elicited key ID93-specific IgA in bronchoalveolar lavage (BAL) samples and polyfunctional Th1-type CD4^+^ T cell numbers in the lungs of BCG-primed mice, at higher levels than in BCG-primed mice followed by i.m. boosts with ID93 + GLA-SE or by i.m. boosts with ID93 + GLA-3M-052-LS. Moreover, boosting BCG-primed mice with ID93 + GLA-3M-052-LS administered by the i.m. route followed by the i.n. route protected mice from weight loss, significantly reduced bacterial burden in the lungs, and reduced inflammatory cells in lung granulomas following low-dose *Mtb* aerosol challenge relative to unimmunized mice, BCG vaccinated mice, and BCG-primed mice boosted only by the i.m. route.

## MATERIALS AND METHODS

### Vaccine formulation composition and manufacturing

ID93 was produced by the Biovac Institute (Cape Town, South Africa) as part of a technology transfer collaboration with the Access to Advanced Health Institute (AAHI, Seattle, WA). Aluminum oxyhydroxide (Alhydrogel 2%) and GLA were acquired from Croda (Princeton, NJ). The 3M-052 was provided by Solventum (Maplewood, MN). Diclofenac sodium salt, poloxamer 188, α-tocopherol, and glycerol were obtained from Spectrum Chemical (New Brunswick, NJ). Squalene was acquired from Sigma (St. Louis, MI). DMPC, DPPC, DOPC, DOTAP, and DSPG were purchased from Lipoid (Ludwigshafen, Germany). DPPE-PEG2000 was acquired from Corden Pharma (Boulder, CO). Plant-derived cholesterol was provided by Wilshire Technologies (Princeton, NJ). Poly(acrylic acid) 5 kDa was purchased from Polysciences (Warrington, PA). Buffer salts were obtained from J. T. Baker (Phillipsburg, NJ).

The adjuvant formulations tested are shown in **Supplementary Table S1**. GLA-SE, 3M-052-AF, NanoAlum, and GLA-3M-052-LS were prepared by high-pressure homogenization essentially as described.^7–10^ The 3M-052-NanoAlum was prepared by mixing 3M-052-AF with NanoAlum and water at a 17:22:61 v:v:v ratio. The 3M-052-Alum was prepared by mixing 3M-052-AF with Alhydrogel 2% and water at a 17:20:63 v:v:v ratio.^11^ Diclofenac liposomes (diclofenac-LS) were prepared by adding diclofenac sodium salt, DOPC, DOTAP, and cholesterol in a 1:7.2:0.8:2 weight ratio to a glass round-bottomed flask, and a chloroform:methanol 2:1 (v:v) ratio was added to solubilize the powders. Following rotary evaporation of the organic solvent overnight, 25 mM ammonium phosphate buffer was added to the flask, which was then placed in a sonicating water bath at 60°C for 1 h. The crude liposomes were then homogenized at 20,000 psi for 5 cycles in an LM20 Microfluidizer (Microfluidics, Westwood, MA) and filtered through a 0.8/0.2-µm polyethersulfone (PES) membrane.

### Adjuvant formulation characterization and stability

Adjuvant formulations were stored at 2-8°C, and physicochemical stability was monitored for 12 months by assessing visual appearance, pH, particle size, and adjuvant concentration. The pH was measured using a Fisherbrand accumet AB150 meter (Fisher Scientific, Ottawa, Canada) with either an Orion ROSS Combination Semi-micro Electrode or an Orion PerpHecT ROSS Combination Micro Electrode (Thermo Fisher Scientific, Waltham, MA). Particle size was measured by dynamic light scattering (Malvern Panalytical Zetasizer, Malvern, UK) essentially as described,^12^ wherein formulations were diluted 1:100 or 1:10 in ultrapure water prior to measurement. Adjuvant concentration was measured by absorbance at 285 nm for diclofenac formulations diluted 1:20 in ethanol, reverse-phase HPLC (Agilent 1100 or 1200, Agilent Technologies, Santa Clara, CA) with charged aerosol detection (Corona Veo, Thermo Fisher Scientific) for GLA, and reverse-phase HPLC with diode array detection (Agilent 1100 or 1200) for 3M-052.^12^ For particle size and HPLC measurements of GLA and 3M-052 content, three to nine replicate measurements of a single sample were conducted. For pH and diclofenac content measurements, a single measurement was conducted.

### Antigen-adjuvant mixing compatibility

Mixtures of the ID93 Ag with each adjuvant formulation were prepared to mimic the compositions employed in the mouse immunization studies described below. The Ag-adjuvant mixtures were monitored at 0, 4, and 24 h after mixing and storage at 2-8°C by assessing visual appearance, pH, particle size, and Ag SDS-PAGE profile. The pH and particle size were measured as described above for the adjuvant formulations. SDS-PAGE was employed to assess Ag integrity and consisted of mixing the sample with 4x loading buffer, heating for 5 min at 90°C, and centrifuging for 30 s at a low speed in a benchtop microfuge. The sample was then loaded onto a Tris-glycine 4-20% gel, run for 65 min at 180 V, and rinsed with water for 5 min before fixing twice for 30 min each with a solution of water containing 50% methanol and 7% acetic acid. The gels were stained overnight using SYPRO Ruby stain (Invitrogen, Carlsbad, CA) and washed with water containing 10% methanol and 7% acetic acid for 30 min followed by rinsing with water 5 times. The gels were imaged on a ChemiDoc MP Imaging System (Bio-Rad Laboratories, Hercules, CA).

### Experimental design for *in vivo* studies

#### Mice

Male and female CC004 mice were acquired from the Systems Genetics Core Facility at the University of North Carolina (UNC).^13^ The CC lines were originally generated and bred at Tel Aviv University in Israel^14^ and Oak Ridge National Laboratory in the US.^15^ Mice were between 6 and 13 weeks of age at study onset. Experiments were conducted under the approved Bloodworks Northwest Research Institute’s Institutional Animal Care and Use Committee (IACUC) protocols #5389-01 and 5389-02 and Texas Biomedical Research Institute’s IACUC protocol #1836 MU. Mice were maintained in microisolator cages, with sterilized water and chow *ad libitum*, in a standard ABSL1/2 vivarium for vaccine immunogenicity studies or ABSL3 vivarium for *Mtb* infections. For all studies, mice were acclimatized after receipt for at least 1 week prior to immunizations. For *Mtb* infections, immunized mice were acclimatized in the ABSL3 facility for at least 1 week prior to challenge studies. Mice were euthanized 4 weeks after *Mtb* infection. No mice met IACUC-approved early removal criteria during either immunization or challenge phases of 20% body weight loss, respiratory distress, or severe lethargy.

#### Experimental rigor

CC004 mice were randomized by cage assignment. For the final three challenge experiments, mice were weighed prior to cage assignments, and cage assignments were made by maintaining approximately equal total mouse weight per study group. Investigators were not blinded to the study groups during data collection and analysis. Quantitative, unbiased assays were developed with SOPs to minimize data biases. Sample sizes were determined based on previous experience in vaccine adjuvant studies regarding the minimum number of mice (*n* = 4 to 6/group/immunogenicity experiment) necessary to identify statistically significant differences between experimental groups.

Negative and positive controls were employed in each study to establish consistency across experiments. Approximately equal numbers of male and female CC004 mice were used in the immunogenicity and challenge studies. Selected readouts were statistically analyzed to determine differences in response based on sex. Four separate protective efficacy experiments were conducted involving the same experimental groups (total *n* = 29 to 40/group). Data from all individual animals were quality assured/quality controlled for technical errors. Prior to final analysis (using data from all four challenge experiments necessary to achieve statistical power), normality and outlier tests were performed. One outlier per group was excluded only if statistically justified using Grubb’s outlier test.

#### Vaccinations and *Mtb* infections

Two immunogenicity-only and four efficacy experiments were performed. Mice received intradermal (i.d.) or subcutaneous (s.c.) BCG priming doses for immunogenicity and challenge experiments, respectively, and i.n. and/or i.m. doses for the test vaccine as indicated. Morbidity was assessed by collecting bodyweights during each experiment. Blood was collected via retro-orbital bleeds performed on isoflurane-sedated mice for immunogenicity experiments. At the end of each experiment, mice were humanely euthanized using CO_2_ as the primary method, and cervical dislocation or vital organ removal as the secondary method. After euthanasia, blood was collected by cardiac puncture, and serum was processed and stored at −20°C or −80°C until use. Harvested tissues were stored on wet ice for transport before processing.

Harvested spleens were gently pressed through a 70-µm mesh cell strainer to create single-cell suspensions and centrifuged at 400 x *g* for 5 min at 4°C to pellet the cells. Red blood cells were lysed with ACK lysis buffer (ThermoFisher) on ice for 30 s, then quenched by adding RPMI medium, and centrifuged again at 400 x *g* for 10 min at 4°C. Supernatants were discarded, and cell pellets were resuspended in RPMI medium. Cell suspensions were filtered through a 2-mL AcroPrep filter plate (Pall Corporation, Port Washington, NY) at 400 x *g* for 5 min at 4°C. Filtered samples were resuspended in RPMI medium containing 10% fetal bovine serum (FBS). Finally, 1-2 x 10^6^ cells/well were transferred to 96-well non-binding U-bottom plates in preparation for stimulation.

Lung tissue was disassociated using a gentleMACS Dissociator (Miltenyi Biotec, Gaithersburg, MD), and single-cell suspensions were created following enzymatic digestion (Hanks’ Balanced Salt Solution supplemented with 10% Liberase [MilliporeSigma], 10% aminoguanidine, 0.1% KN-62, and 1.25% DNase). Lung cell samples were incubated at 37°C and 5% CO_2_ for 30 min. Samples were processed on the gentleMACS Dissociator, using the m_lung_02.01 program. Lung cells were then washed with RPMI medium, filtered through a 70-µm MACS SmartStrainer (Miltenyi Biotec), counted on a Guava easyCyte cytometer (Cytek Biosciences, Fremont, CA), and then plated in 96-well round-bottom plates at 1 x 10^6^ cells per well in RPMI medium + 10% FBS in preparation for stimulation.

#### ID93 stimulation and flow cytometry

Cells were stimulated in RPMI medium containing 10% FBS, 50 μM 2-mercaptoethanol, α-CD28 (BD Biosciences), and one of three stimulations: 0.26% DMSO or phosphate-buffered saline (PBS) as a negative control; 1 μg/well (10 μg/mL) of ID93 in PBS; or 10 μg/well of PMA-ionomycin solution as a positive control. Plates were incubated for 2 h at 37°C and 5% CO_2_, after which brefeldin A (BioLegend) was added, and plates were incubated for an additional 6 h at 37°C and 5% CO_2_.

After incubation, plates were centrifuged at 400 x *g* for 3 min, supernatants were removed, and cells were washed twice with PBS. Viability was assessed using Zombie Green (BioLegend), and Fc receptors were blocked with CD16/CD32 antibody (Ab) (Invitrogen). Cells were surface stained for mouse CD4 (PerCPCy5.5 or APC-Cy7, BD Biosciences), CD8 (BV510, BD Biosciences), and CD44 (APC-Cy7 or PE-CF594, BD Biosciences). Spleen cells were additionally stained for mouse CD107a (APC, BioLegend). Lung cells were additionally stained for mouse CD154 (BV605, BD Biosciences), CD69 (PE, BD Biosciences), and CD103 (BV711, BioLegend). After extracellular staining, cells were fixed and permeabilized using BD Cytofix/Cytoperm (BD Biosciences) and stained for intracellular cytokines with mouse TNFα (BV421, BioLegend), IL-2 (PE-Cy5, BioLegend), IFNγ (PE-Cy7, BD Biosciences), IL-5 (PE or APC, BioLegend), and IL-17A (AF700, BD Biosciences). Spleen cells were additionally stained for mouse IL-10 (BV711, BD Biosciences). Cells were run on an LSR Fortessa (BD Biosciences) or CytoFLEX (Beckton Dickson) flow cytometer and analyzed with FlowJo v.10 (BD Biosciences). Representative gating strategies are shown in **Supplementary Figures S1-S2**.

Background signal from unstimulated wells was subtracted from the ID93-stimulated signal for each mouse, and resulting negative values were assigned a value of zero.

#### Serum ID93-specific IgG, IgG1, and IgG2a; BAL and lung IgA by ELISA

Detection of serum ID93-specific IgG, IgG1, and IgG2c was performed as previously described.^16^ For immunogenicity experiments, 384-well flat-bottom polystyrene high-binding plates (Corning, Corning, NY) were coated with 1 µg/mL of ID93, produced and purified in house, and incubated overnight at 4°C. CC004 naïve mouse serum was used as a negative control. A custom anti-ID93-Rv3619 IgG Ab (Antibody Solutions, Santa Clara, CA) was used as a positive control for total IgG quantification. Serum anti-ID93 Abs were detected using an Anti-Mouse IgG (Fc Specific)-Alkaline Phosphatase Ab (Sigma-Aldrich #A2429), Anti-Mouse IgG1 (γ-chain Specific)-Alkaline Phosphatase Ab (Sigma-Aldrich #SAB3701172), and Anti-Mouse IgG2a (γ-chain Specific)-Alkaline Phosphatase Ab (Sigma-Aldrich #SAB3701179). Detection of BAL IgA was performed as previously described.^17^ Briefly, plates were coated as above, and each BAL sample was diluted 1:4 before 1:2 serial dilution in a non-binding 384-well plate (Corning #3684) to generate a 14-point dilution curve. BAL anti-ID93 Abs were detected using an Anti-Mouse IgA (α-chain specific)-Alkaline Phosphatase Ab (Sigma-Aldrich #A4937). Plates were developed using phosphatase substrate tablets (Sigma-Aldrich #S0942) dissolved in diethanolamine substrate buffer (Fisher Scientific #PI34064). Plates were read using a spectrophotometer at 405 nm after a 30-min incubation in the dark. Absolute quantification of serum IgG titers were interpolated from the linear region of each sample dilution curve using a 4-point logistic curve fit of the positive control dilution curve.

Endpoint BAL IgA, serum IgG1, and serum IgG2c titers were calculated by performing a least-squares fit of OD values to a 4-point sigmoidal curve. For *Mtb* challenge studies, serum ID93-specific IgG and lung homogenate ID93-specific IgA assays used the same methods modified for 96-well format.

#### Bone marrow ELISpot assay

Plates were coated with recombinantly expressed ID93. Single-cell suspensions of bone marrow samples were collected during necropsy, seeded at 1.0 x 10^6^ cells per well, subsequently serially diluted, and then probed with a goat anti-mouse IgG horseradish peroxidase conjugate Ab (SouthernBiotech #1036-05). A 3-amino-9-ethylcarbazole (AEC) substrate kit was used as the colorimetric substrate for the ELISpot development. The plates were incubated overnight in the dark before the reaction was quenched. The resulting colored spots were enumerated using an automated ELISpot reader (CTL Analyzer, Cellular Technology Limited, Cleveland, OH). Data were analyzed using ImmunoSpot software (Cellular Technology Limited).

#### Desirability index methodology

To generate a meaningful overall ranking of the adjuvant formulations tested, we employed a desirability function approach adapted from previous reports.^12,18,19^ Briefly, data from multiple readouts were log-transformed and the median was calculated among all animals in the same group, and the resulting values were normalized on a unitless scale of 0 to 1. In one study, only lung T cell intracellular cytokine staining data were available due to spleen samples being compromised during sample handling. A weighted composite desirability score (*D*) was then calculated using the equation:

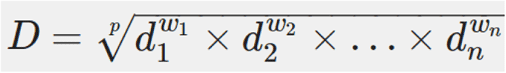

where *d_i_* = partial desirability attributed to the *i*th immunological response (*i* = 1, 2, …, *n*), *w_i_* = weighting attributed to the *i*th response, and 𝑝𝑝 = ∑^𝑛𝑛^ 𝑤𝑤_𝑖𝑖_. The weighting system was designed to rank the various immunological readouts, with 5 representing of greatest importance and 1 representing of least importance. For example, Th1 and Th17 CD4 T cell responses in the lung are considered important for protection against TB and were thus given a high weight (see **Tables S2 and S3** for all assigned weight parameters).

#### *Mtb* aerosol challenge and bacterial lung burden

Mice were exposed to an aerosol of virulent *Mtb* strain Erdman using an inhalation exposure system (Glas-col, Terre Haute, IN) calibrated to deliver approximately 25-100 colony forming units (CFUs) to the lungs of each individual mouse as described.^20–22^ Four weeks post challenge, lung *Mtb* burden was calculated by plating serial dilutions of whole lung homogenates onto oleic acid-albumin-dextrose-catalase (OADC) enrichment supplemented 7H11 agar plates. *Mtb* CFUs were counted after 4 weeks at 37°C and transformed to a log_10_ scale. Mice were checked daily for routine health observations and weighed at least twice weekly post infection.

#### Histology and automated image analysis

After euthanasia, one lung lobe from each challenge study mouse was inflated and fixed in 10% neutral buffered formalin, processed, embedded in paraffin, sectioned at 5 microns thick, stained with hematoxylin and eosin (H&E) and digitally scanned at 40x magnification using an Olympus scanner. Granulomas in the lung sections and individual cell types (macrophages, lymphocytes, and neutrophils, based on morphology) were detected and segmented from background by using a board-certified veterinary pathologist’s (G.B.) manual training annotations in the Aiforia Create (v.6.0) platform with default parameter settings (Aiforia Technologies, Helsinki, Finland). Iterative rounds of training, testing, and validation were performed until errors were minimized for granuloma and cell identification, *i.e.*, false positive rates 1.77% and 0.59%; and false negative rates 0.67% and 0.39%, respectively.

#### Statistical analyses

Immunogenicity and protective efficacy data were analyzed by one-way ANOVA (α = 0.05) with Dunnett’s, Brown-Forsythe and Welch ANOVA (α = 0.05) with Dunnett’s T3, or the non-parametric Kruskal-Wallis test (α = 0.05) with Dunn’s correction for multiple comparisons. For immunogenicity studies, hypothesis testing included each experimental group compared to the ID93 + GLA-SE immunized group and by route of administration for select studies. For challenge studies, hypothesis testing included BCG-primed groups compared to the unimmunized group and comparison between all BCG-primed groups (GraphPad Prism v. 10.0.2 software). Ab, ELISpot, and CFU data were log-transformed prior to analysis. Statistical significance was determined by *p* value of <0.05. Ab measurements below the limit of detection (LOD) were assigned an arbitrary low value of 1. To evaluate the impact of sex on immunogenicity responses, data were analyzed by two-way ANOVA with Sidak’s correction for multiple comparisons or unpaired *t-*test pairwise comparisons.

## RESULTS

### Adjuvant formulations demonstrate long-term physicochemical stability

We sought to improve vaccine-specific local respiratory and systemic immune responses to the clinical-stage TB Ag ID93 by pairing it with a panel of adjuvant formulations with different mechanisms of action, as well as exploring the immunological effects of different immunization routes. Adjuvant formulations and doses were selected from recently developed and published work developing novel adjuvant compositions for other indications or were newly developed and reported here for the first time (**Supplementary Table S1**). Each adjuvant formulation was characterized for visual appearance, pH, particle size, and adjuvant content following manufacture. Adjuvant formulations remained physicochemically stable for at least 16 to 24 months stored at 2-8°C (**Figure 1A-D**) including no notable changes in visual appearance. To determine physicochemical compatibility of the ID93 Ag and the adjuvant formulations after mixing, we assessed the same parameters as above as well as ID93 Ag degradation by SDS-PAGE at 0, 4, and 24 h after mixing. No detrimental impacts on ID93 Ag or the adjuvant formulations were apparent after mixing (**Figure 1E-H**). Having demonstrated acceptable physicochemical stability and compatibility, all of the Ag-adjuvant vaccine combinations were determined to be suitable for advancement into the mouse model for immunogenicity evaluation.

**Figure 1.**
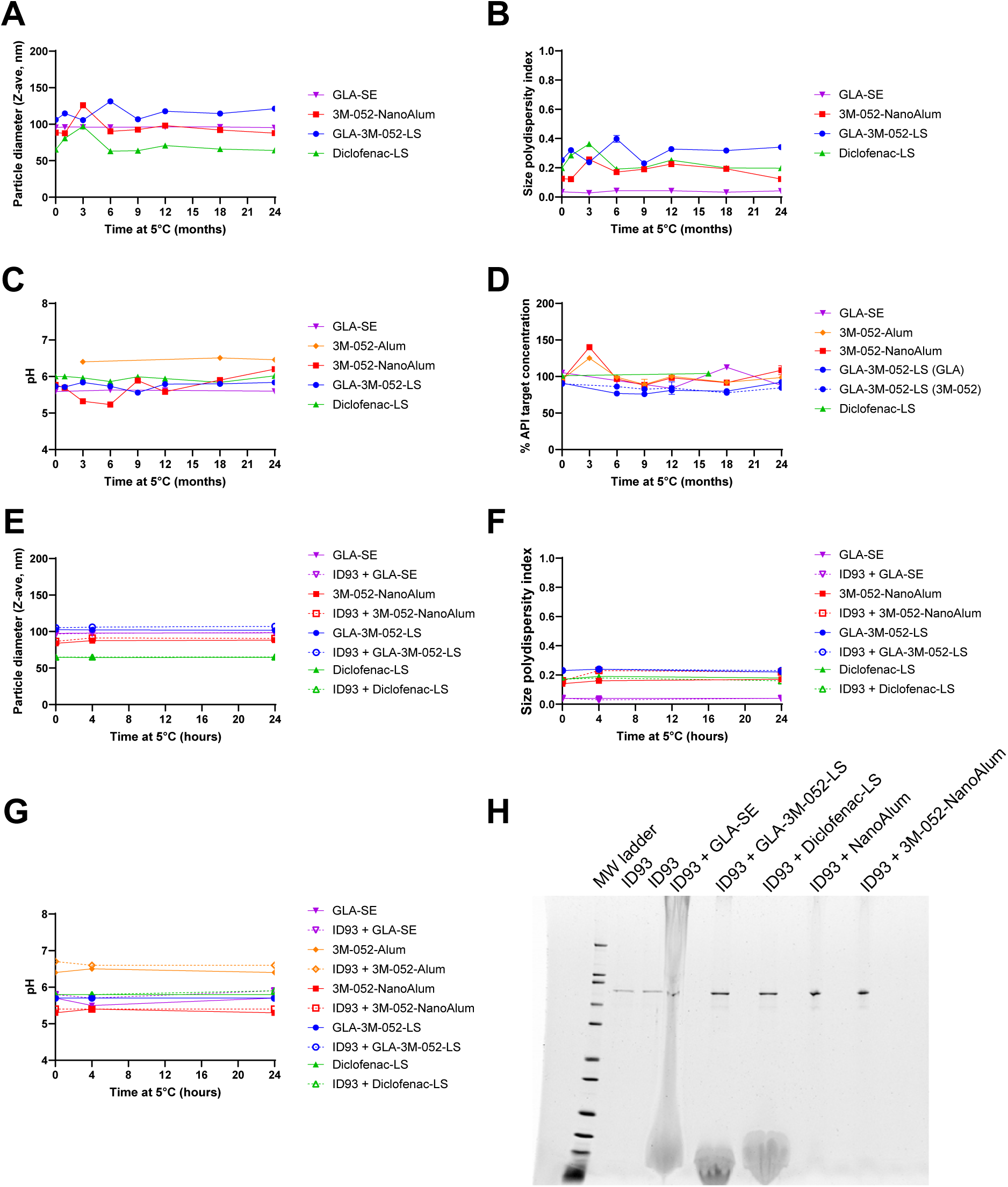
Adjuvant formulation stability and mixing compatibility with ID93 antigen. Long-term adjuvant formulation stability at 5°C was assessed by measuring **(A)** particle diameter, **(B)** size polydispersity index, **(C)** pH, and **(D)** active pharmaceutical ingredient (API) content. Short-term mixing compatibility of adjuvant formulations with ID93 at 5°C were assessed by **(E)** particle diameter, **(F)** size polydispersity index, **(G)** pH, and **(H)** SDS-PAGE with fluorescent stain at 4 h after mixing with the indicated formulations. 3M-052-Alum is not included in the particle size or SDS-PAGE measurements because the large, heterogeneous particles in the Alum-based formulation are outside of the range of the dynamic light scattering technique and introduces artifacts into the SDS-PAGE gels.

### Immunogenicity studies

To determine the effects of the adjuvant formulations on ID93-specific systemic and mucosal immune responses, experimental groups of CC004 mice (*n* = 4-6/group) were immunized twice, 4 weeks apart, according to the regimens described in **Table 1**. Immunized CC004 mice showed no signs of overt toxicity based on appearance or behavior. Average weight loss of ≤5% within 2 days following immunization was observed in some groups, after which weight was generally maintained or increased (**Supplementary Figure S3**), consistent with previously reported data.^12^ Three weeks after the first immunization, serum samples were collected to assess Ab responses. At 1 week following the second immunization, mice were necropsied, and serum, BAL, bone marrow, spleen, and lung tissue samples were collected to assess Ab and cellular responses.

**Table 1.** Experimental group and regimen description for adjuvant screening immunogenicity study.

| Experimental Group# | Immunization #1<br>(Day 0) | Immunization #2<br>(Day 28) |
| --- | --- | --- |
| 1 | ID93 + saline (i.m.) | ID93 + saline (i.m.) |
| 2 | ID93 + GLA-SE (i.m.) | ID93 + GLA-SE (i.m.) |
| 3 | ID93 + 3M-052-Alum (i.m.) | ID93 + 3M-052-Alum (i.m.) |
| 4 | ID93 + 3M-052-NanoAlum<br>(i.m.) | ID93 + 3M-052-NanoAlum<br>(i.m.) |
| 5 | ID93 + GLA-3M-052-LS (i.m.) | ID93 + GLA-3M-052-LS (i.m.) |
| 6 | ID93 + GLA-3M-052-LS (i.n.) | ID93 + GLA-3M-052-LS (i.n.) |
| 7 | ID93 + Diclofenac-LS (i.m.) | ID93 + Diclofenac-LS (i.m.) |

Each adjuvant formulation and immunization route induced a distinct immune response profile. For example, the highest levels of ID93-specific serum IgG titers at 3 weeks following the first immunization were elicited by 3M-052-Alum (i.m.) and 3M-052-NanoAlum (i.m.), whereas the GLA-3M-052-LS (i.n.) and Diclofenac-LS (i.m.) groups showed no benefit in serum IgG stimulation relative to ID93 (i.m.) alone (**Supplementary Figure S4**). However, at 1 week following the second immunization, the GLA-3M-052-LS (i.m.) group showed the highest serum IgG titers and the greatest number of long-lived Ab-secreting cells in the bone marrow, whereas the GLA-3M-052-LS (i.n.) group showed the greatest median magnitude serum IgG2a/IgG1 ratio although the variation in response between mice in this group was also high (**Figure 2A-C**). Mucosal IgA in BAL fluid was most evident in the GLA-3M-052-LS (i.n.) and 3M-052-NanoAlum (i.m.) groups (**Figure 2D**).

**Figure 2.**
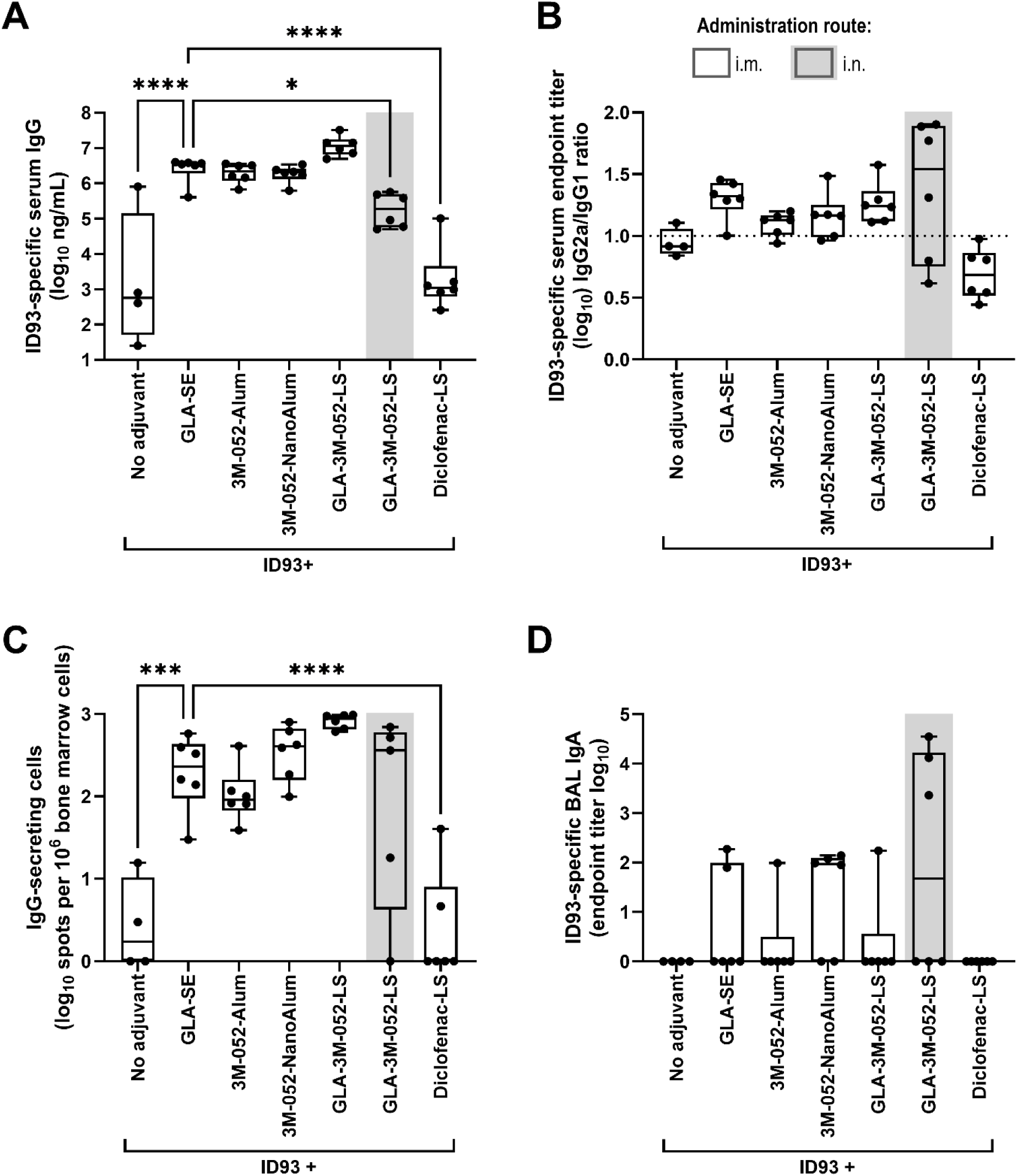
Post-boost antibody immunogenicity elicited by homologous vaccine regimens involving ID93 with distinct adjuvant formulations. CC004 mice (*n* = 4 to 6/group) were immunized according to the regimens described in Table 1. One week after the second immunization, ID93-specific immune responses were measured in the serum, bone marrow, and bronchoalveolar lavage (BAL) fluid. **(A)** Serum IgG (see Supplementary Figure S4 for post-prime data), **(B)** serum IgG2a/IgG1 ratio, **(C)** long-lived IgG-secreting cells in the bone marrow, **(D)** BAL IgA. **(A, C)** Data were log-transformed and analyzed using one-way ANOVA with Dunnett’s multiple comparisons test. **(D)** Due to the small group size, the statistical analysis was carried out in the most conservative manner possible, employing the non-parametric Kruskal-Wallis test with Dunn’s correction for multiple comparisons. \**p* < 0.05, *** *p* < 0.001, **** *p* < 0.0001. Bars indicate median values, boxes indicate the 25 to 75% spread, and whiskers indicate the minimum and maximum values, with individual data points shown.

For cellular immune responses, GLA-3M-052-LS (i.m.) elicited high percentages of cytokine-producing CD4^+^ T cells in the spleen, comparable to the responses elicited by the GLA-SE benchmark control group (**Figure 3A**), and 3M-052-NanoAlum also induced notable levels of cytokine-producing CD4^+^ T cells in the spleen. In both cases, a Th1 quality response was apparent as indicated by IFNγ, TNFα, and polyfunctional T cells. Interestingly, cytokine-producing CD4^+^ T cells in the lung elicited by the adjuvant-containing formulations were minimal and not higher than the levels achieved with Ag alone (**Figure 3B)**. Data for each mouse, plots, and statistical analysis for each cytokine are also reported in **Supplementary Figures S5-S6** for CD4^+^ and CD8^+^ T cells in the spleen and lung. Two-way ANOVA analysis of the impact of sex on selected immune readouts indicated sex contributed to variation for some readouts, although the percent of total variation attributable to sex was less than that attributable to immunization regimen (2.0% to 9.3% *vs*. 30.5% to 81.2%).

**Figure 3.**
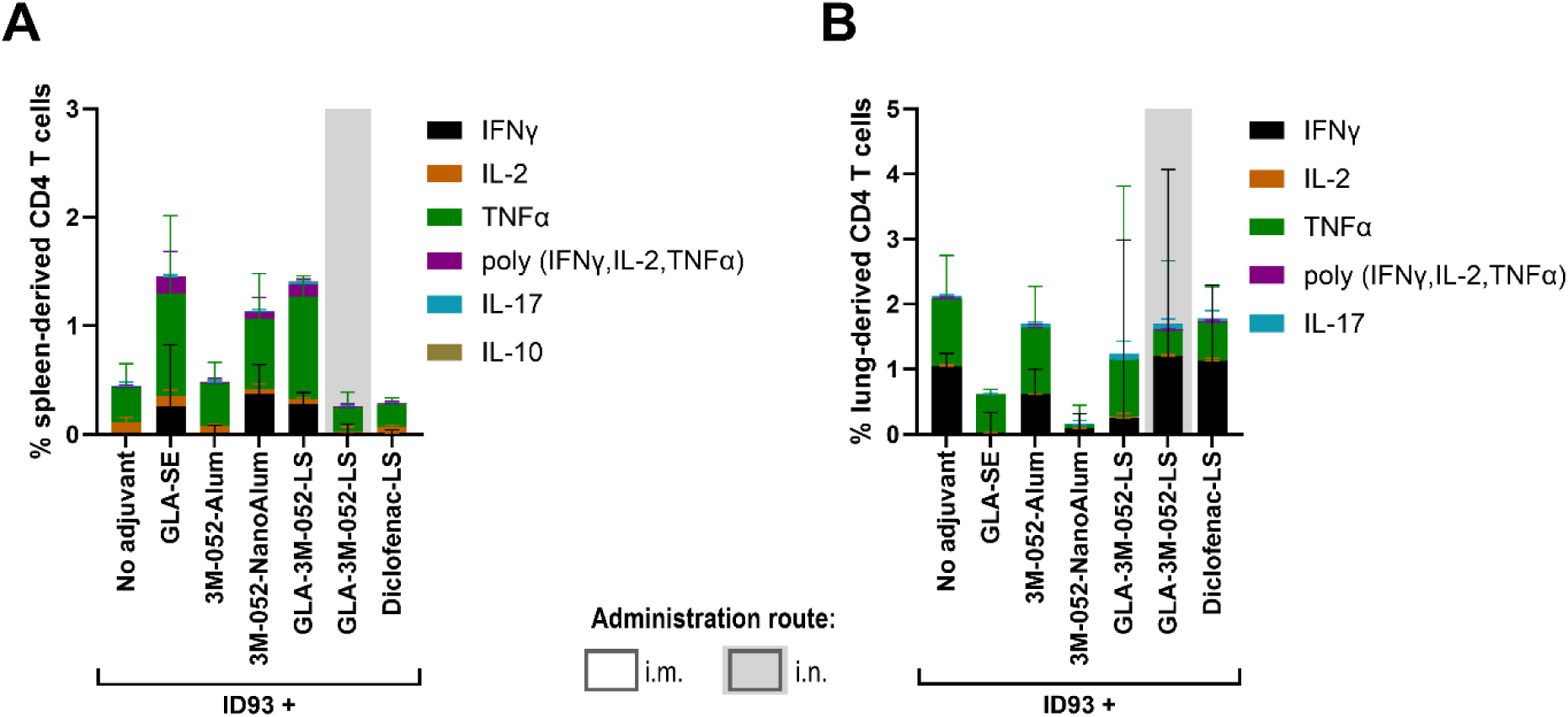
Cellular immunogenicity elicited by homologous vaccine regimens involving ID93 with distinct adjuvant formulations. CC004 mice (*n* = 4 to 6/group) were immunized according to the regimens described in Table 1. One week after the second immunization, ID93-specific immune responses were measured in the spleen and lungs. **(A)** Percent cytokine-producing CD4^+^ T cells in the spleen, **(B)** percent cytokine-producing CD4^+^ T cells in the lung. Bars indicate median values, and error bars indicate the interquartile spread. Statistical analysis was not conducted on the stacked data shown here but was conducted on individual cytokine data as represented in Supplementary Figures S5 and S6.

To rank overall immune response profiles, a desirability index approach was used to incorporate the post-boost immune responses into a single score, weighted according to perceived importance of each immune measure to protective efficacy or immune response durability (**Supplementary Table S2**). Highest weights were assigned to mucosal immune responses, such as Th17 cellular immunity, Th1 cellular immunity in the lung, and long-lived Ab-secreting cells. In this context, the desirability index score indicated that GLA-3M-052-LS (i.n.) and GLA-3M-052-LS (i.m.) induced the highest quality overall immune response, substantially higher than the GLA-SE (i.m.) control adjuvant (**Table 2**). The desirability index for 3M-052-NanoAlum (i.m.) was slightly higher than that of the GLA-SE (i.m.) control adjuvant, while 3M-052-Alum (i.m.) and Diclofenac-LS (i.m.) did not improve the desirability index score compared to ID93 Ag alone, and were indicated to be inferior to the control GLA-SE (i.m.) adjuvant.

**Table 2.** Desirability index ranking for adjuvant screening immunogenicity study.

| Experimental Group | Desirability Index Score |
| --- | --- |
| ID93 + GLA-3M-052-LS (i.n.) | 0.26 |
| ID93 + GLA-3M-052-LS (i.m.) | 0.24 |
| ID93 + 3M-052-NanoAlum (i.m.) | 0.19 |
| ID93 + GLA-SE (i.m.) | 0.13 |
| ID93 (i.m.) | 0.11 |
| ID93 + 3M-052-Alum (i.m.) | 0.10 |
| ID93 + Diclofenac-LS (i.m.) | 0.07 |

Based on their performance in immunogenicity studies, GLA-3M-052-LS and 3M-052-NanoAlum were selected for further evaluation in CC004 mice (*n* = 5 to 6/group) previously vaccinated with BCG to mimic real-use scenarios for these new vaccine candidates. Various combinations of i.m. immunization routes for the 3M-052-NanoAlum adjuvant and i.m. and i.n. immunization routes for the GLA-3M-052-LS adjuvant were administered (**Table 3**). GLA-SE (i.m.) was again used as a benchmark control adjuvant, and BCG alone was used as an unboosted control. The adjuvant-containing boost formulations were administered 28 and 56 days after a BCG priming. Four weeks after the final immunization, serum, BAL, bone marrow, and lung tissue samples were collected to assess Ab and cellular responses.

**Table 3.** Experimental group and regimen description for lead candidate immunogenicity study.

| Experimental Group# | Immunization #1<br>(Day 0) | Immunization #2<br>(Day 28) | Immunization #3<br>(Day 56) |
| --- | --- | --- | --- |
| 1 | BCG Pasteur (i.d.) | Saline (i.m.) | Saline (i.m.) |
| 2 | BCG Pasteur (i.d.) | ID93 + GLA-SE (i.m.) | ID93 + GLA-SE (i.m.) |
| 3 | BCG Pasteur (i.d.) | ID93 + 3M-052-NanoAlum<br>(i.m.) | ID93 + 3M-052-NanoAlum<br>(i.m.) |
| 4 | BCG Pasteur (i.d.) | ID93 + 3M-052-NanoAlum<br>(i.m.) | ID93 + GLA-3M-052-LS (i.n.) |
| 5 | BCG Pasteur (i.d.) | ID93 + GLA-3M-052-LS (i.m.) | ID93 + GLA-3M-052-LS (i.m.) |
| 6 | BCG Pasteur (i.d.) | ID93 + GLA-3M-052-LS (i.m.) | ID93 + GLA-3M-052-LS (i.n.) |
| 7 | BCG Pasteur (i.d.) | ID93 + GLA-3M-052-LS (i.n.) | ID93 + GLA-3M-052-LS (i.m.) |
| 8 | BCG Pasteur (i.d.) | ID93 + GLA-3M-052-LS (i.n.) | ID93 + GLA-3M-052-LS (i.n.) |

Levels of ID93-specific serum IgG and IgG-secreting long-lived plasma cells elicited by i.m. boost vaccinations of ID93 + 3M-052-NanoAlum or ID93 + GLA-3M-052-LS were similar to those elicited by ID93 + GLA-SE (**Figure 4A-B**). Interestingly, the groups of mice immunized with a heterologous i.m.-i.n. or homologous i.n.-i.n. ID93 + GLA-3M-052-LS boost regimen retained similar levels of IgG-secreting long-lived plasma cells as the ID93 + GLA-SE group while also showing strong Th1-type CD4^+^ T cell responses in the lung and statistically significant mucosal IgA compared to the ID93 + GLA-SE group (**Figure 4B-D**). Data from each mouse, plots, and statistical analysis for each cytokine are also reported in **Supplementary Figure S7** for CD4^+^ and CD8^+^ T cells in the lung. Two-way ANOVA analysis of the impact of sex on selected immune readouts indicated it was a source of variation for some readouts, although the percent of total variation attributable to sex was much lower than that attributable to immunization regimen (1.8% to 3.4% *vs*. 67.4% to 72.9%).

**Figure 4.**
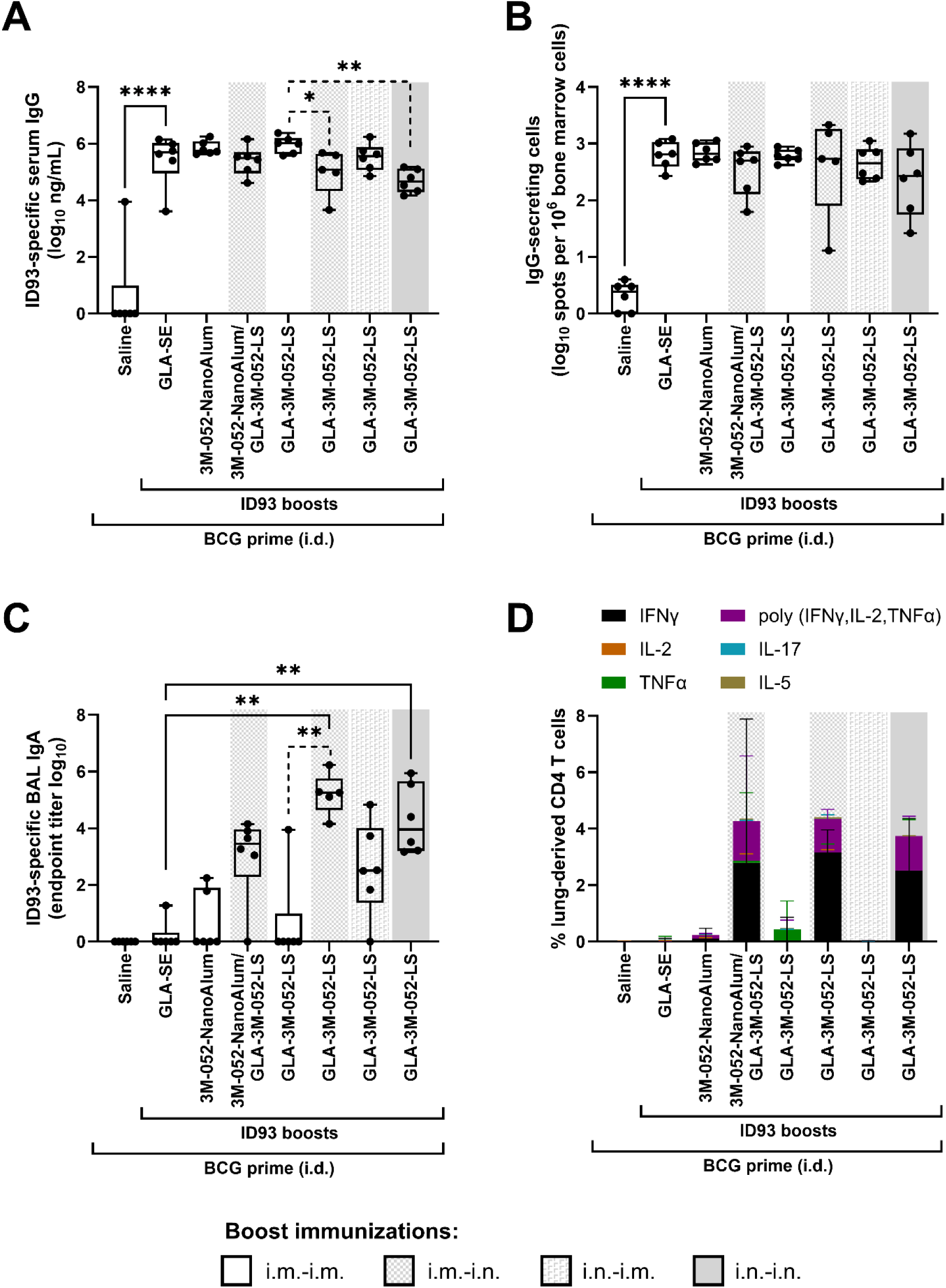
Immunogenicity elicited by heterologous vaccine regimens involving BCG prime and ID93 boosts with lead candidate adjuvant formulations. CC004 mice (*n* = 5-6/group) were immunized according to the regimens described in Table 3. Four weeks after the final immunization, ID93-specific immune responses were measured in the serum, bone marrow, bronchoalveolar lavage (BAL) fluid, and lung. **(A)** Serum IgG, **(B)** long-lived IgG-secreting cells in the bone marrow, **(C)** mucosal IgA, **(D)** percent cytokine-producing CD4^+^ T cells in the lung. **(A-B)** Data were log-transformed and analyzed using one-way ANOVA with Dunnett’s multiple comparisons test. **(C)** Due to the small group size, the statistical analysis was carried out in the most conservative manner possible, employing the non-parametric Kruskal-Wallis test with Dunn’s correction for multiple comparisons. Solid lines: each experimental group compared to the ID93 + GLA-SE immunized group; dotted lines: comparison between all routes of administration for ID93 + GLA-3M-052-LS boosts. * *p* < 0.05, ** *p* < 0.01, **** *p* < 0.0001. Bars indicate median values, boxes indicate the 25-75% spread, and whiskers indicate the minimum and maximum values, with individual data points shown. **(D)** Statistical analysis was not conducted on the stacked data shown but was conducted on individual cytokine data as represented in Supplementary Figure S7.

Consistent with results from the previous experiment, the adjuvant compositions (including GLA-SE) administered only by the i.m. route did not induce substantial mucosal immunity, and the adjuvanted vaccine administered only by the i.n. route resulted in the lowest magnitude of systemic antibody response although not statistically distinct from ID93 + GLA-SE (**Figure 4**). The desirability index score for each vaccine regimen, weighted according to perceived importance of selected immune responses (**Supplementary Table S3**), was highest with heterologous GLA-3M-052-LS (i.m.-i.n.) boosting, followed by heterologous 3M-052-NanoAlum (i.m.)/GLA-3M-052-LS (i.n.) boosting and homologous GLA-3M-052-LS (i.n.) boosting (**Table 4**). Thus, the ID93 + GLA-3M-052-LS (i.m.-i.n.) heterologous immunization regimen appeared to achieve the most comprehensive immune response profile including maintaining a high level of systemic immunity stimulation while also eliciting substantial mucosal immune responses.

**Table 4.** Desirability index ranking for lead candidate immunogenicity study.

| Experimental Group (all groups BCG-primed except for the last group) | Desirability Index Score |
| --- | --- |
| ID93 + GLA-3M-052-LS (i.m.-i.n.) | 0.29 |
| ID93 + 3M-052-NanoAlum (i.m.)/GLA-3M-052-LS (i.n.) | 0.21 |
| ID93 + GLA-3M-052-LS (i.n.-i.n.) | 0.16 |
| ID93 + 3M-052-NanoAlum (i.m.-i.m.) | 0.09 |
| ID93 + GLA-3M-052-LS (i.n.-i.m.) | 0.06 |
| ID93 + GLA-3M-052-LS (i.m.-i.m.) | 0.05 |
| ID93 + GLA-SE (i.m.-i.m.) | 0.03 |
| Saline (i.m.-i.m.) | 0.02 |
| Saline without BCG prime (i.m.-i.m.) | 0.01 |

### Protective efficacy: Reduction in lung *Mtb* burden

The lead vaccine candidate was then tested in four independent efficacy experiments conducted following the experimental design in **Table 5 and Figure 5A**. Briefly, CC004 mice (total *n* = 29 to 40/group) were primed with BCG, rested for 6 weeks, and then immunized twice, 3 weeks apart with the lead vaccine candidate, ID93 + GLA-3M-052-LS, delivered either in i.m.-i.m. or i.m.-i.n. route regimens. Three weeks following the final immunization, mice were challenged with a low dose (25-100 CFUs) of aerosolized *Mtb* Erdman. Four weeks after *Mtb* infection, lungs and blood were harvested to assess *Mtb* lung burden and ID93-specific Abs. Combined data from the four experiments are shown as a reduction in log_10_ CFUs compared to the mean CFUs in unimmunized mice challenged with *Mtb* (**Figure 5B**). CFU data from individual experiments and other control groups are shown in **Supplementary Figure S8**. All challenge data were analyzed for sex-specific responses, and none were detected (**Supplementary Figure S9**).

**Figure 5.**
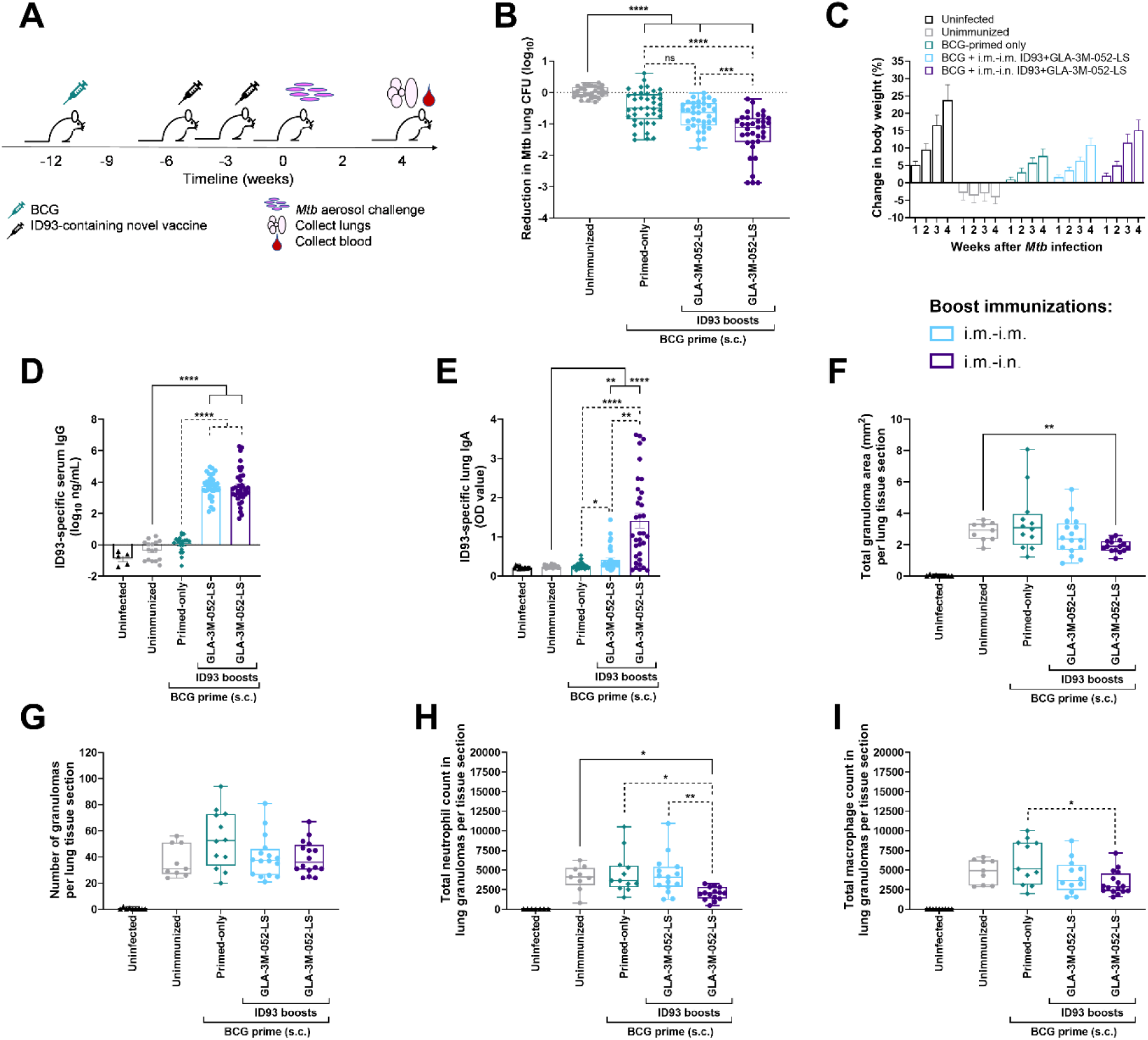
Protective efficacy elicited by heterologous vaccine regimens in BCG-primed and ID93 + GLA-3M-052-LS immunized CC004 mice. **(A)** Experimental design for *Mtb* challenge studies in BCG-primed mice. Three weeks after the final immunization, mice were challenged with a low dose (25-100 CFUs) of aerosolized *Mtb* strain Erdman. Four weeks later, serum and lungs were harvested. **(B)** Lung CFU data were log-transformed and plotted as reduction in CFUs (log_10_) compared to the average of unimmunized mice infected with *Mtb*. **(C)** Change in body weight after *Mtb* infection, compared to baseline immediately prior to infection. **(D)** ID93-specific IgG in serum. **(E)** ID93-specific IgA in lung. **(F)** Total area of granuloma per lung tissue section. **(G)** Number of granulomas per lung tissue section.**(H)** Number of neutrophils in granulomas per lung tissue section. **(I)** Number of macrophages in granulomas per lung tissue section. Box-whisker plots show the median and interquartile range with whiskers indicating the minimum and maximum values and individual data points shown. **(B-E)** show combined data (total *n* = 29 to 40 CC004 mice/group) from four separate experiments (shown in Supplementary Figure S8) immunized according to the regimens described in Table 5. **(F-I)** show one representative experiment where there were sufficient granulomas to analyze. Statistical analysis was conducted using **(B,D,F-I)** Brown-Forsythe and Welch ANOVA with Dunnett’s T3 or **(E)** the non-parametric Kruskal-Wallis test with Dunn’s correction for multiple comparisons (solid lines: BCG-primed groups compared to unimmunized group; dotted lines: comparison between all BCG-primed groups); * *p* < 0.05, ** *p* < 0.01, *** *p* < 0.001, **** *p* < 0.0001.

**Table 5.**
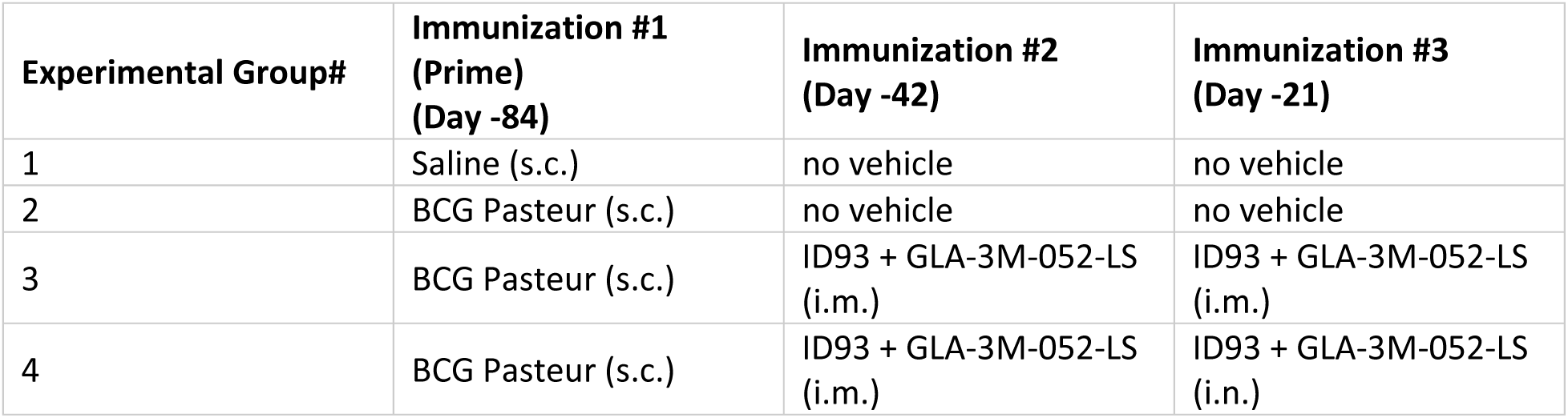
Experimental group and regimen description for protective efficacy studies.

All BCG-vaccinated groups demonstrated a significant reduction in lung CFUs compared to unimmunized mice (**Figure 5B**). This reduction was notably further reduced by heterologous i.m.-i.n. boosting with ID93 + GLA-3M-052-LS. In contrast, animals immunized with ID93 + GLA-3M-052-LS by the homologous i.m.-i.m. regimen did not achieve greater protection than animals immunized with BCG alone. Thus, boosting BCG with a heterologous regimen of an adjuvanted subunit vaccine candidate (ID93 + GLA-3M-052-LS) administered by the i.m. route followed by the i.n. route resulted in improved protective efficacy in controlling *Mtb* burden in the lungs than immunization with BCG alone.

### Protective efficacy: Improved ID93-specific responses and weight maintenance after challenge correspond to reduced lung inflammation

We also evaluated the impact of vaccination on post-challenge body weight; ID93-specific serum IgG and lung IgA; and granuloma number and area, and numbers of inflammatory cells in lungs of vaccinated then *Mtb-*challenged male and female CC004 inbred mice. All vaccines protected mice from weight loss following *Mtb* infection, and improved weight gain nearly comparable to age-matched uninfected mice (**Figure 5C**). Boosting BCG-primed mice with ID93 + GLA-3M-052-LS either i.m.-i.m. or i.m.-i.n induced significantly higher levels of post-challenge ID93-specific serum IgG compared to both unimmunized and BCG vaccinated mice (**Figure 5D**), suggesting the two boost vaccine regimens elicit similar levels of systemic humoral immunity. Notably, the heterologous i.m.-i.n boosting with ID93 + GLA-3M-052-LS had the largest positive effect on ID93-specific lung IgA present after challenge, significantly increased compared to all other groups including i.m.-i.m. boosted mice (**Figure 5E**), suggesting that mucosal immunization uniquely results in higher mucosal immune responses after challenge. By histopathology evaluation and image analysis, we observed a trend for i.m.-i.n. heterologous boost to reduce the total granuloma area per lung tissue section, which reached statistical significance when compared to unimmunized mice (**Figure 5F**) without significantly affecting the number of granulomas per lung tissue section (**Figure 5G**). The numbers of neutrophils in lung granulomas, which are generally considered indicators of non-protective/detrimental innate inflammation and disease progression in primary *Mtb* infection,^23,24^ were significantly reduced by i.m.-i.n. heterologous boost compared to other vaccine groups (**Figure 5H**). The number of macrophages in lung granulomas was less impacted by vaccination although there was a trend for BCG vaccination to increase macrophages compared to unvaccinated mice, an increase that was prevented by both i.m.-i.m and i.m.-i.n. boosting with ID93 + GLA-3M-052-LS, with statistical significance achieved with the heterologous i.m.-i.n. boost vaccination regimen (**Figure 5I**).

## DISCUSSION

By systematically comparing Ab and cellular immune responses induced by the ID93 Ag paired with different adjuvant systems and immunization routes, we identified ID93 + GLA-3M-052-LS administered by a heterologous i.m.-i.n. route regimen as a promising new TB vaccine candidate that elicits comprehensive mucosal and systemic immunity and protects CC004 inbred mice, a strain reported to be susceptible to *Mtb* and poorly protected by BCG vaccination,^6^ from low-dose aerosolized *Mtb* challenge. This study advances important concepts in TB vaccine development, including evaluation of new adjuvant formulations and mucosal immunization routes as potentially key for development of more effective next-generation vaccines. Additionally, this work establishes the use of an *Mtb*-susceptible CC mouse strain for assessment of the immunogenicity and efficacy of clinically relevant TB vaccine compositions to model vulnerable individuals who may not be protected against pulmonary TB by BCG vaccination.

The GLA-3M-052-LS adjuvant system administered s.c., i.m., or i.n. has been successfully employed with recombinant *Entamoeba histolytica* and SARS-CoV-2 vaccine candidates in mouse and non-human primate models to elicit mucosal and systemic immune responses.^10,12,25^ However, this is the first evaluation of GLA-3M-052-LS in a TB vaccine candidate, as well as the first time this adjuvant formulation is administered using a heterologous i.m.-i.n. route immunization regimen.

Comprised of clinical-stage synthetic TLR ligands and lipid excipients demonstrating robust physicochemical stability and compatibility with different routes of delivery, GLA-3M-052-LS represents a promising adjuvant system for next-generation TB vaccine candidates. Although ID93 + GLA-3M-052-LS administered by a homologous i.m.-i.m. regimen was a strong performer in terms of immunogenicity, it did not provide additional protection from *Mtb* challenge in BCG-primed mice compared to BCG alone. Thus, the heterologous route regimen of ID93 + GLA-3M-052-LS in BCG-primed mice was necessary to improve upon the protective efficacy induced by BCG alone. Studies of other vaccine candidates have indicated that a heterologous i.m.-i.n. immunization regimen, also known as a ‘prime and pull’ strategy, represents a promising approach for eliciting systemic and mucosal immune responses.^26,27^ The 3M-052-NanoAlum formulation is another notable adjuvant system that performed well in the immunogenicity studies and is reported here for the first time, although it was only tested by the i.m. route as we considered the presence of aluminum nanoparticles inappropriate for i.n. administration.

CC recombinant inbred mouse strains were derived from a multi-parental mouse reference population. Individual CC strains are increasingly being used to study how pathogens interact with new mouse genetic backgrounds^28,29^ and are being used in panels to represent genetic diversity that is more similar to the human population.^28^ Here, we used CC004 inbred mice as an immune competent model for immunogenicity and *Mtb* challenge studies because the strain is susceptible to *Mtb* and less protected by BCG vaccination than the standard C57BL/6 inbred mouse strain.^6^ These phenotypes make the CC004 inbred strain an attractive preclinical animal model of vulnerable humans who may not be fully protected against pulmonary TB by BCG vaccination. Following low-dose *Mtb* challenge within the range of the infectious dose for humans,^30^ we observed that the novel vaccine regimen of ID93 + GLA-3M-052-LS administered by heterologous boost (i.m.-i.n.) provided the best protection against *Mtb,* noted by greatest reduction in bacterial load. This corresponded with the strongest post-challenge systemic humoral (serum ID93-specific IgG) and mucosal (lung ID93-specific IgA) immune responses. Heterologous i.m.-i.n. boosting of prior BCG-priming additionally reduced the total area occupied by granulomas in lung tissue sections and the numbers of innate inflammatory cells (neutrophils and macrophages) within granulomas, suggesting that this novel vaccine regimen provides additional benefits by reducing recruitment of innate inflammatory cells that may contribute to lung tissue damage in pulmonary TB.

We acknowledge a few limitations of the current study, including that the vaccine formulations and *Mtb* challenge studies were tested in one preclinical animal species: CC004 inbred mice. Although the results we obtained are quite promising, using larger and different animal models, such as guinea pigs and non-human primates, could provide additional confidence in meaningful translation to humans, particularly with regards to alternative routes of immunization. Another limitation was that only one CC mouse strain was studied here. Using additional CC strains to validate vaccine immunogenicity and efficacy across different genetic backgrounds known to respond differently to BCG vaccination and *Mtb* challenge would also increase confidence that i.m.-i.n. boosting of BCG with ID93 + GLA-3M-052-LS could induce protection across many genotypes. In order to model the current standard of practice for humans, we focused on BCG priming administered by injection through the skin (i.d. and s.c.) rather than focusing on BCG administered as a booster or by alternative dosing routes. Moreover, we did not deplete specific cell populations, perform adoptive cell transfers, or manipulate Ab responses to elucidate possible mechanisms of protection; this will be considered in future studies. It would be also desirable to establish the protective efficacy of the heterologous i.m.-i.n. ID93 + GLA-3M-052-LS candidate compared to the same candidate administered by the homologous i.n.-i.n. route. Finally, long-term survival studies were not conducted and would be interesting to do in the future. A strength of the current study is that immunogenicity and efficacy was assessed in BCG vaccinated mice, which represents the most relevant real-world use scenario.

Overall, we have demonstrated a systematic approach to next-generation TB vaccine development using CC004 inbred mice to evaluate immunogenicity and efficacy of combinations of new adjuvant systems and immunization routes with the clinical-stage ID93 Ag. The ID93 + GLA-3M-052-LS vaccine formulation administered intramuscularly and intranasally induced robust mucosal and systemic immunity that significantly protected CC004 mice from *Mtb* challenge. The ID93 + GLA-3M-052-LS vaccine candidate thus merits subsequent evaluation in additional animal models, including other CC mouse strains, Diversity Outbred mice, guinea pigs, and non-human primates in preparation for potential clinical testing in the future.

## DATA AVAILABILITY

The datasets generated and/or analyzed during the current study are available from the corresponding author on reasonable request.

## MATERIAL AVAILABILITY

AAHI’s novel adjuvant formulations are available for research and/or clinical use, subject to third party rights for certain indications or territories, under the terms and conditions of an appropriate material transfer agreement (for preclinical research) or clinical supply agreement (for clinical studies). Correspondence and material requests should be addressed to the corresponding author.

## ACKNOWLEDGMENTS

The authors are grateful to Robert Kinsey, Joseph McCollum, Sierra Reed, Jeffrey Guderian, Eduard Melief, Ravi Iyer, Ethan Lo, Sam Beaver, Noah Cross, and Peter Battisti for excellent technical assistance; to Elise Echefu for project management; to Mark Tomai from Solventum for providing 3M-052; to Valerie Soza for scientific editing; and to Texas Biomedical Research Institute’s BSL3 Operations Program for their support.

## FUNDING

Research reported in this publication was supported by the National Institutes of Health (NIH) under the National Institute of Allergy and Infectious Diseases (NIAID), award number NIH/NIAID R61 AI169026. The total project costs (100%) was financed with federal funds. The content is solely the responsibility of the authors and does not necessarily represent the official views of the NIH .NIH/NIAID had no role in study design, data collection, analysis, interpretation, writing, and decision to submit paper for publication.

## AUTHOR CONTRIBUTIONS

**Emily A. Voigt:** Conceptualization, Formal analysis, Resources, Writing – original draft, Writing – review & editing. **Anas Alsharaydeh:** Investigation, Writing – review & editing. **Darshan N. Kasal:** Investigation, Formal analysis, Writing – original draft, Writing – review & editing. **Madeleine F. Jennewein:** Conceptualization, Investigation, Writing – review & editing. **Devin S. Brandt:** Investigation, Formal analysis, Writing – original draft, Writing – review & editing. **Susan Lin:** Conceptualization, Investigation, Writing – review & editing. **Jasneet Singh:** Investigation, Writing – review & editing. **Julie Bakken:** Investigation, Writing – review & editing. **Raodoh Mohamath:** Investigation, Writing – review & editing. **Pauline Fusco:** Investigation, Writing – review & editing. **Jordi B. Torrelles:** Conceptualization, Formal analysis, Resources, Writing – original draft, Writing – review & editing. **Gillian Beamer:** Conceptualization, Formal analysis, Resources, Writing – original draft, Writing – review & editing. **Christopher B. Fox:** Conceptualization, Formal analysis, Resources, Writing – original draft, Writing – review & editing.

## COMPETING INTERESTS

The authors declare no Competing Non-Financial Interests but the following Competing Financial Interests. CBF is an inventor on patents and/or patent applications involving formulations of GLA, NanoAlum, and 3M-052. EAV, JBT, GB, and CBF are co-inventors on US provisional patent application no. 63/781,868 titled “Next-generation Mycobacterium Tuberculosis Vaccine.” All other authors declare that they have no competing interests.

