## Supplementary Material for "The ID93 + GLA-3M-052-LS vaccine candidate elicits mucosal and systemic immunogenicity and protective efficacy against *Mycobacterium tuberculosis* challenge in BCG-primed Collaborative Cross inbred mice"

### Supplementary Tables

#### Table S1. Adjuvant formulation panel.

| Adjuvant Name (Dose) | Description | Immunization Route | Reported Immune Response Profile and References |
|---|---|---|---|
| GLA-SE (5 µg GLA, 2% v/v squalene) | Synthetic TLR4 agonist in squalene oil-in-water emulsion | i.m. | Serum Abs, systemic Th1 $CD4^+$ T cells [1–3] |
| 3M-052-Alum (1 µg 3M-052, 100 µg aluminum) | Synthetic TLR7/8 agonist adsorbed to aluminum oxyhydroxide microparticles | i.m. | Serum Abs, long-lived Ab-secreting cells, systemic Th1 $CD4^+$ T cells [4–6] |
| 3M-052-NanoAlum (1 µg 3M-052, 100 µg aluminum) | Synthetic TLR7/8 agonist adsorbed to aluminum oxyhydroxide nanoparticles stabilized w/poly(acrylic) acid | i.m. | New adjuvant formulation |
| GLA-3M-052-LS (10 µg GLA, 4 µg 3M-052) | Synthetic TLR4 and TLR7/8 agonists in PEGylated liposomes | i.m.or i.n. | Serum and mucosal Abs, long-lived Ab-secreting cells, systemic and mucosal Th1 and Th17 cells [7–9] |
| Diclofenac-LS (50 µg diclofenac) | Mucosal-associated invariant T cell activator formulated with cationic liposomes | i.m. | New adjuvant formulation |

**Table S2. Weighting criteria of immune response readouts for desirability index scoring for the adjuvant screening immunogenicity study.**

| Readout | Weight | Function | Justification |
|---|---|---|---|
| IFNγ$^+$ CD4$^+$ T cells in spleen | 3 | Maximize | Systemic cellular Th1 immunity |
| IFNγ$^+$ CD4$^+$ T cells in lung | 4 | Maximize | Lung cellular Th1 immunity |
| TNFα$^+$ CD4$^+$ T cells in spleen | 2 | Maximize | Systemic cellular Th1 immunity |
| TNFα$^+$ CD4$^+$ T cells in lung | 3 | Maximize | Lung cellular Th1 immunity |
| IL-2$^+$ CD4$^+$ T cells in spleen | 1 | Maximize | Systemic cellular Th1 immunity |
| IL-2$^+$ CD4$^+$ T cells in lung | 2 | Maximize | Lung cellular Th1 immunity |
| IL-17$^+$ CD4$^+$ T cells in spleen | 4 | Maximize | Systemic cellular Th17 immunity |
| IL-17$^+$ CD4$^+$ T cells in lung | 5 | Maximize | Lung cellular Th17 immunity |
| IFNγ$^+$/TNFα$^+$/IL-2$^+$ CD4$^+$ T cells in spleen | 3 | Maximize | Systemic cellular Th1 immunity |
| IFNγ$^+$/TNFα$^+$/IL-2$^+$ CD4$^+$ T cells in lung | 4 | Maximize | Lung cellular Th1 immunity |
| IL-10$^+$ CD4$^+$ T cells in spleen | 1 | Minimize | Undesirable Th2 immune response indicator |
| IFNγ$^+$ CD8$^+$ T cells in spleen | 2 | Maximize | Effector CD8$^+$ T cells in spleen |
| TNFα$^+$ CD8$^+$ T cells in spleen | 1 | Maximize | Effector CD8$^+$ T cells in spleen |
| TNFα$^+$ CD8$^+$ T cells in lung | 2 | Maximize | Effector CD8$^+$ T cells in lung |
| Serum IgG | 4 | Maximize | Serum IgG Ab titers are indicative of systemic immunogenicity |
| IgG2/IgG1 ratio | 4 | Maximize | IgG2c/IgG1 ratio correlates with Th1 immunity |
| IgA in BAL | 5 | Maximize | Mucosal IgA Ab titers are indicative of lung immunogenicity |
| IgG-secreting cells in bone marrow | 4 | Maximize | Long-lived plasma cells are indicator of durable Ab-mediated immunity |

**Table S3. Weighting criteria of immune response readouts for desirability index scoring for the lead candidate immunogenicity study.**

| Readout | Weight | Function | Justification |
|---|---|---|---|
| $IFN\gamma^+$ $CD4^+$ T cells in lung | 4 | Maximize | Lung cellular Th1 immunity |
| $TNF\alpha^+$ $CD4^+$ T cells in lung | 3 | Maximize | Lung cellular Th1 immunity |
| $IL\text{-}17^+$ $CD4^+$ T cells in lung | 5 | Maximize | Lung cellular Th17 immunity |
| $IFN\gamma^+/TNF\alpha^+/IL\text{-}2^+$ $CD4^+$ T cells in lung | 4 | Maximize | Lung cellular Th1 immunity |
| $IL\text{-}5^+$ $CD4^+$ T cells in lung | 1 | Minimize | Undesirable Th2 immune response indicator |
| $IFN\gamma^+$ $CD8^+$ T cells in lung | 3 | Maximize | Effector $CD8^+$ T cells in lung |
| $TNF\alpha^+$ $CD8^+$ T cells in lung | 2 | Maximize | Effector $CD8^+$ T cells in lung |
| Serum IgG | 4 | Maximize | Serum IgG Ab titers are indicative of systemic immunogenicity |
| IgA in BAL | 5 | Maximize | Mucosal IgA Ab titers are indicative of lung immunogenicity |
| IgG-secreting cells in bone marrow | 4 | Maximize | Long-lived plasma cells are indicator of durable Ab-mediated immunity |

**Supplementary Figures**

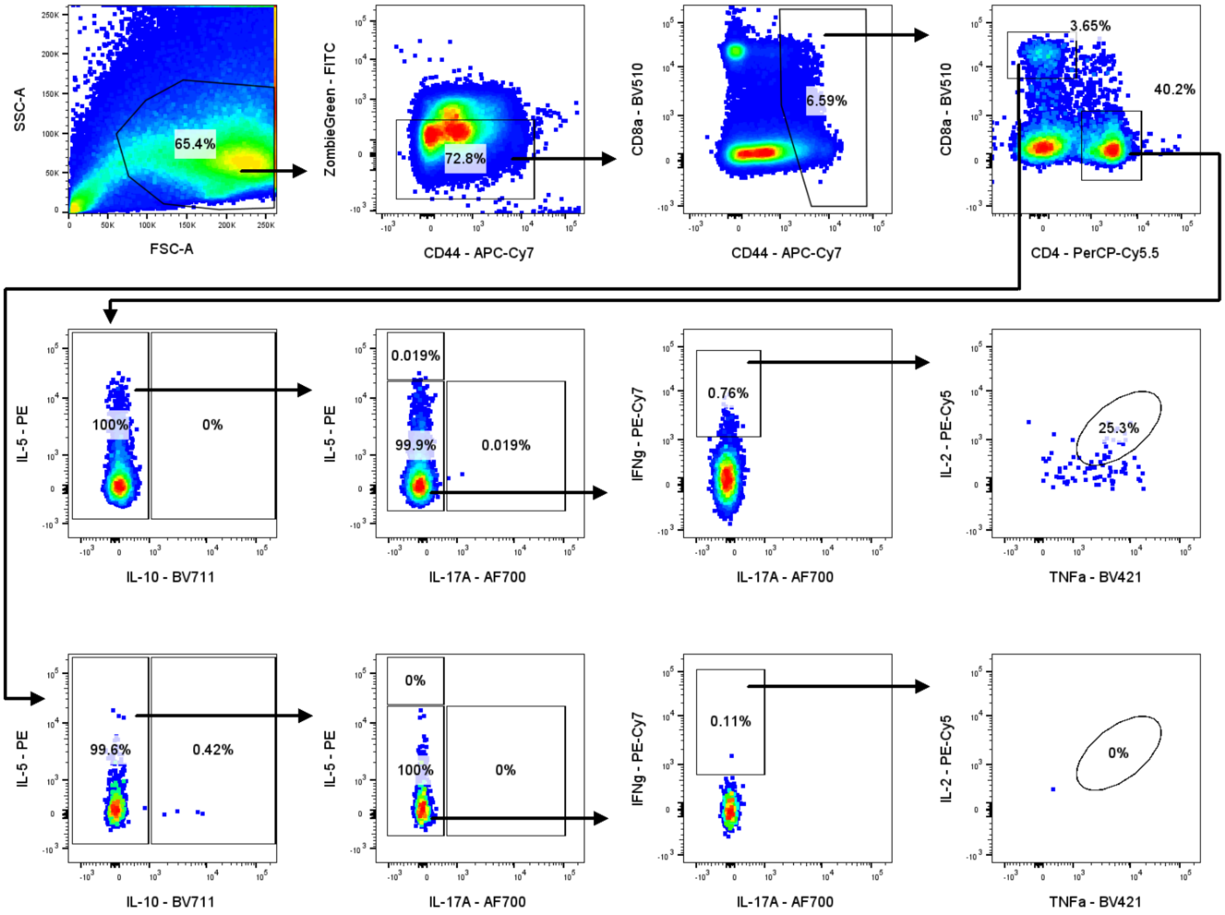


**Figure S1. Spleen flow cytometry gating strategy.** Splenocytes were stained and analyzed via flow cytometry. Cells were first gated on total lymphocytes followed by live cells ($FITC^-$) and then activated cells ($CD44^+$). Individual gates were drawn for $CD4^+$ and $CD8^+$ T cells. Within each of these subsets, specific cytokines were gated using biplots, including IL-10, IL-5, IL-17A, IFNγ, IL-2, and TNFα.

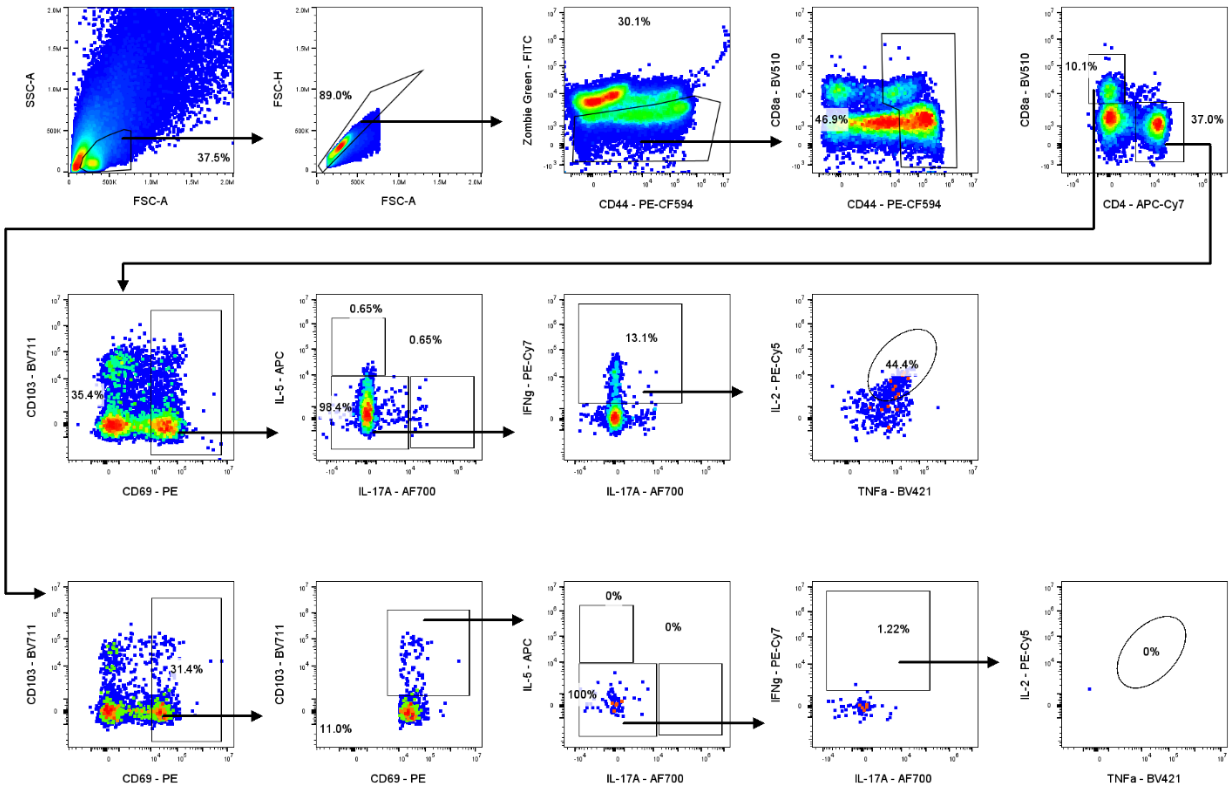


**Figure S2. Lung flow cytometry gating strategy.** Lung cells were stained and analyzed via flow cytometry. Cells were first gated on total lymphocytes followed by live cells (FITC$^-$) and then activated cells (CD44$^+$). Individual gates were drawn for CD4$^+$ and CD8$^+$ T cells, and further gated for tissue-resident markers (CD69$^+$ for both and CD103$^+$ for CD8$^+$ T cells). Within each of these subsets, specific cytokines were gated using biplots, including IL-5, IL-17A, IFNγ, IL-2, and TNFα.

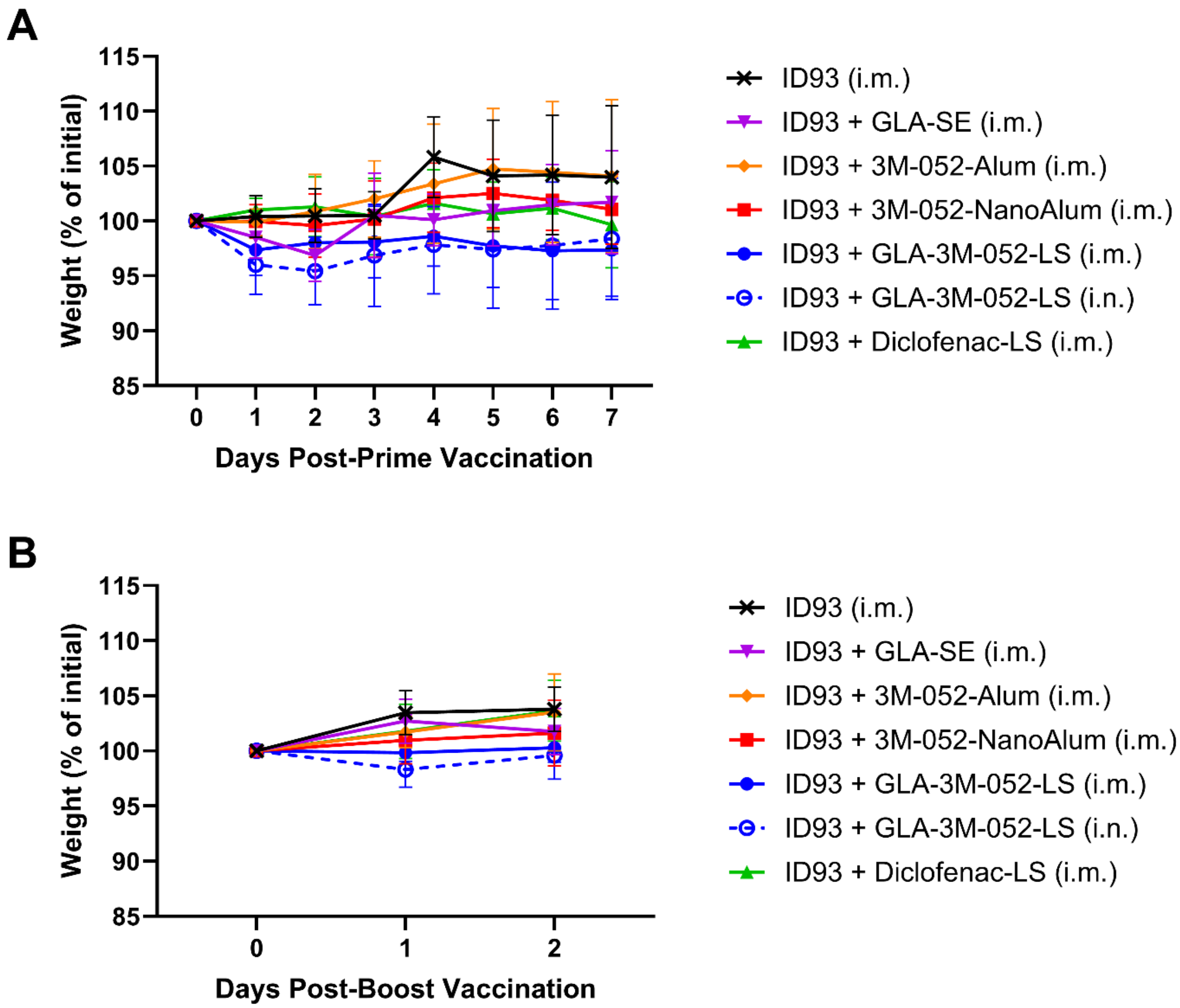


**Figure S3. Mouse weight following immunization.** Change in weight following **(A)** prime immunization and **(B)** boost immunization with the indicated vaccine formulations (*n* = 4-6/group).

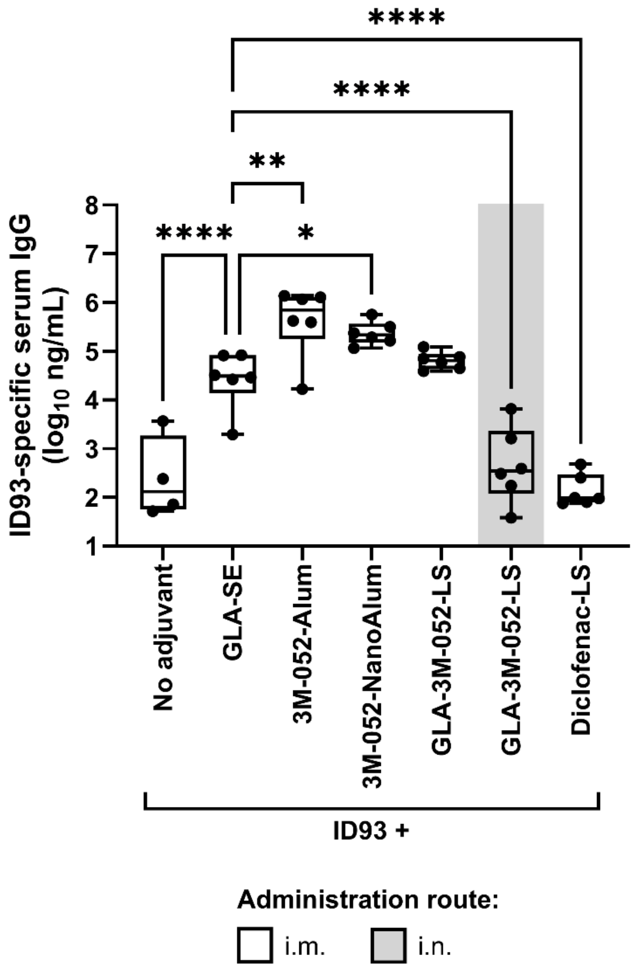


**Figure S4. Post-prime antibody immunogenicity elicited by vaccine regimens involving ID93 with distinct adjuvant formulations.** CC004 mice ($n$ = 4 to 6/group) were immunized according to the regimens described in Table 1. Three weeks after the first immunization, ID93-specific IgG was measured in the serum. Data were log-transformed and analyzed using one-way ANOVA with Dunnett's multiple comparisons test; *$p < 0.05$, ** $p < 0.01$, **** $p < 0.0001$. Bars indicate median values, boxes indicate the 25-75% spread, and whiskers indicate the minimum and maximum values, with individual data points shown.

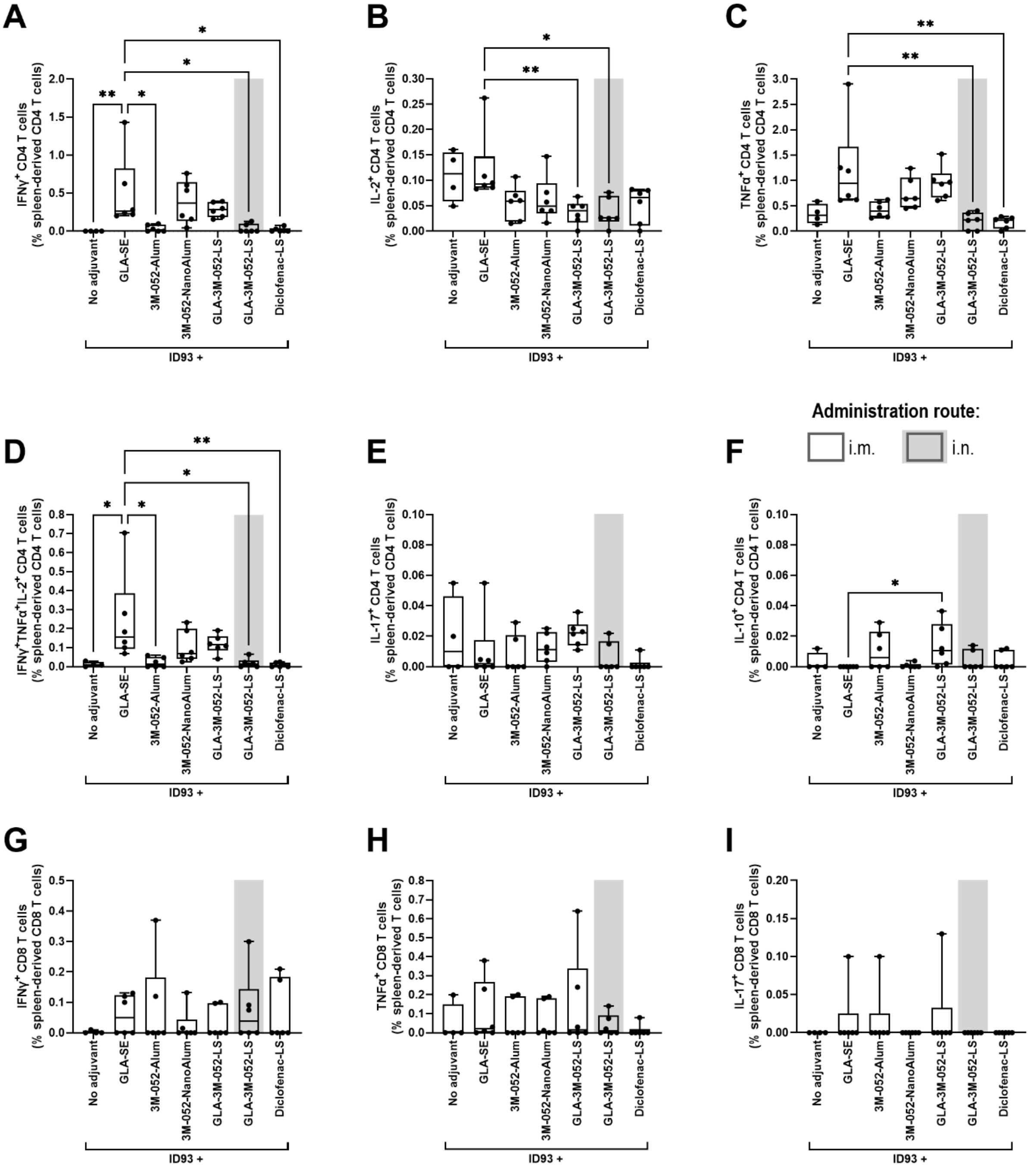


**Figure S5. Cytokines elicited in the spleen by CD4⁺ or CD8⁺ T cells by homologous vaccine regimens involving ID93 with distinct adjuvant formulations.** CC004 mice ($n$ = 4 to 6/group) were immunized according to the regimens described in Table 1. One week after the second immunization, ID93-specific immune responses were measured in the spleen. Due to the small group size, the statistical analysis was carried out in the most conservative manner possible, employing the non-parametric Kruskal-Wallis test with Dunn's correction for multiple comparisons; * $p < 0.05$, ** $p < 0.01$, *** $p < 0.001$. Bars indicate median values, boxes indicate the 25-75% spread, and whiskers indicate the minimum and maximum values, with individual data points shown.

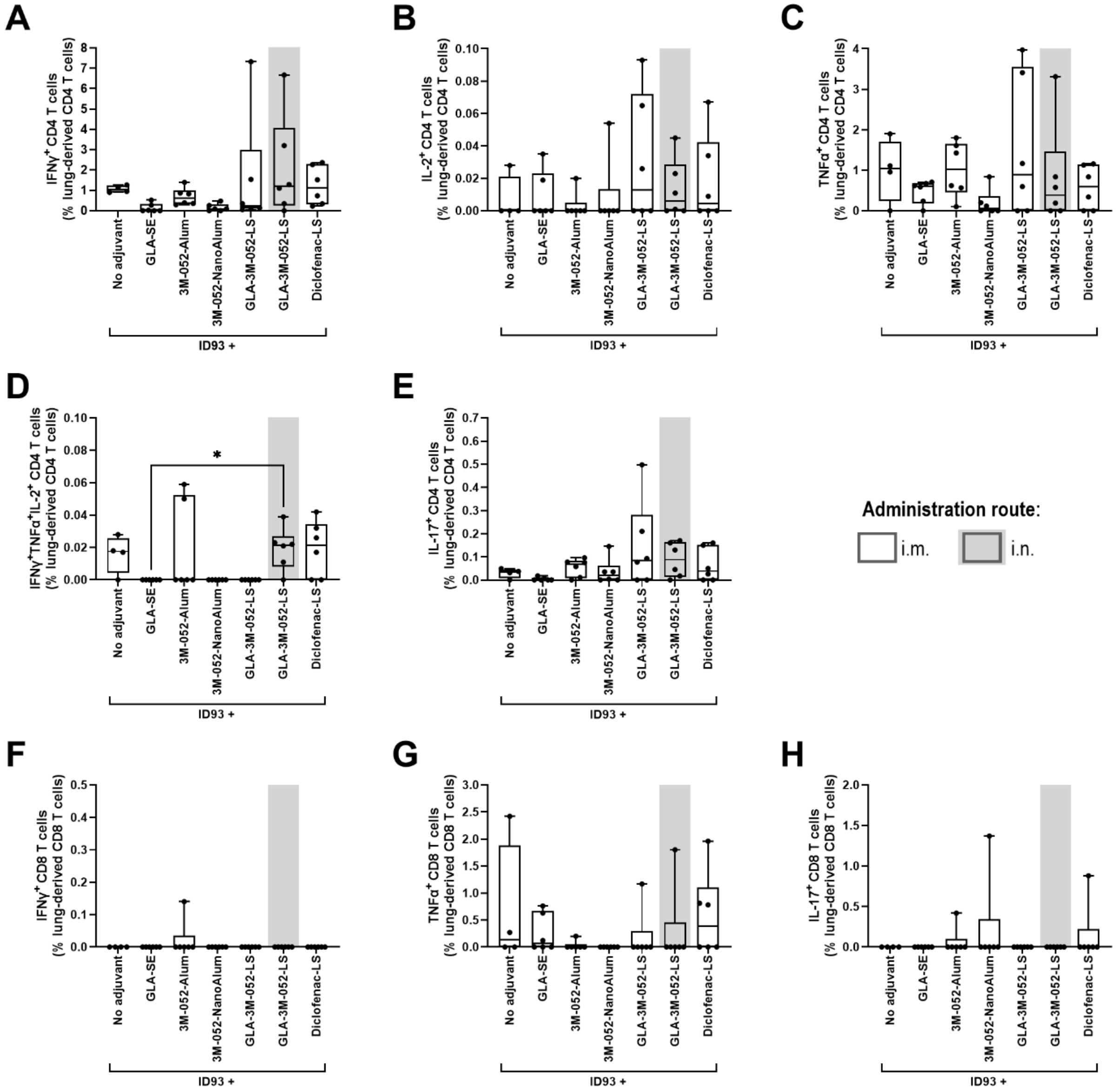


**Figure S6. Cytokines elicited in the lung by CD4+ or CD8+ T cells by homologous vaccine regimens involving ID93 with distinct adjuvant formulations.** CC004 mice ($n$ = 4 to 6/group) were immunized according to the regimens described in Table 1. One week after the second immunization, ID93-specific immune responses were measured in the lung. Due to the small group size, the statistical analysis was carried out in the most conservative manner possible, employing the non-parametric Kruskal-Wallis test with Dunn's correction for multiple comparisons; * $p < 0.05$, ** $p < 0.01$, *** $p < 0.001$. Bars indicate median values, boxes indicate the 25 to 75% spread, and whiskers indicate the minimum and maximum values, with individual data points shown.

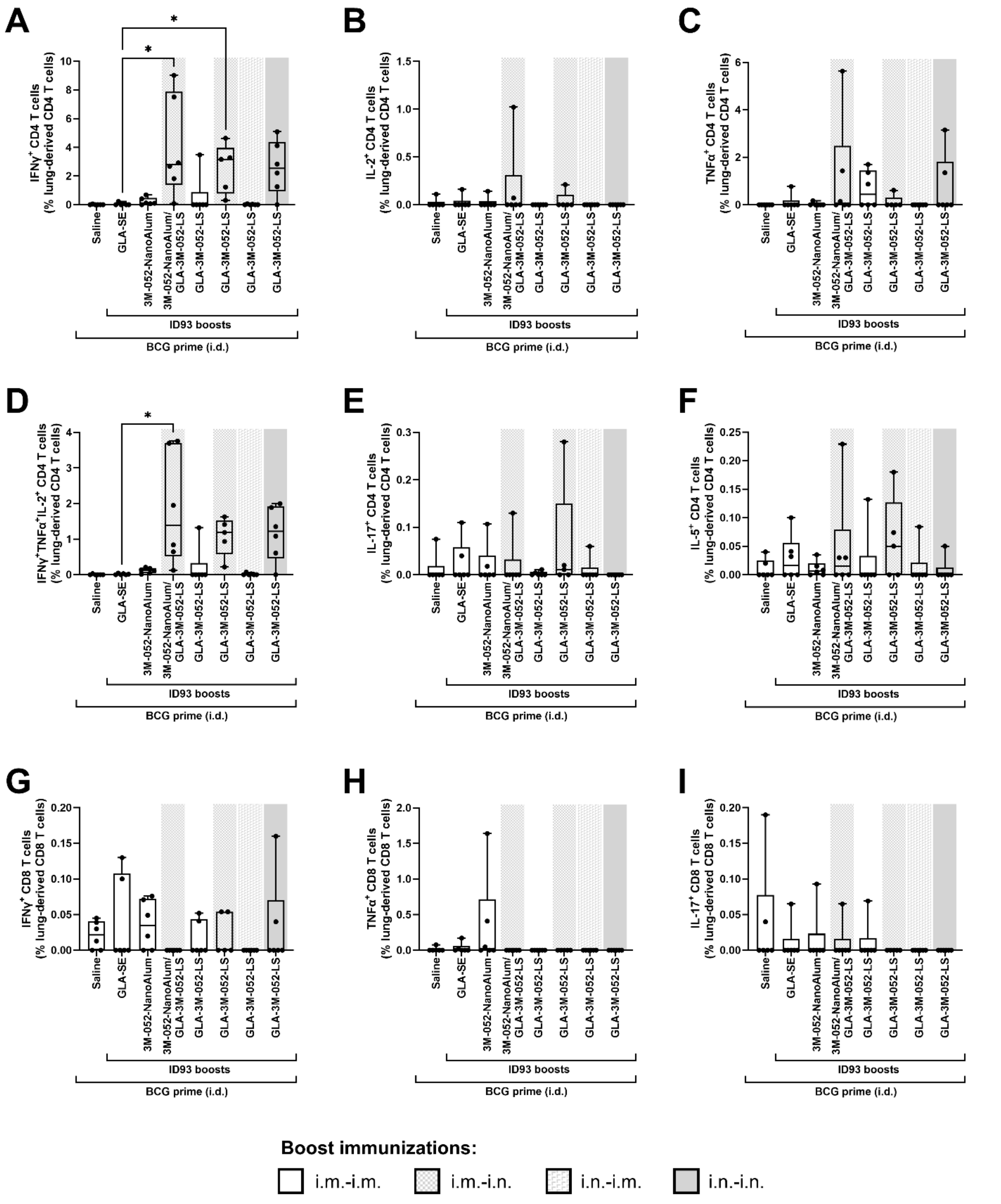


**Figure S7. Cytokines elicited in the lung by CD4+ or CD8+ T cells by heterologous vaccine regimens involving BCG prime followed by intramuscular or intranasal ID93 boosts with distinct adjuvant formulations.** CC004 mice ($n$ = 5 to 6/group) were immunized according to the regimens described in Table 3. Four weeks after the final immunization, ID93-specific immune responses were measured in the lung. Due to the small group size, the statistical analysis was carried out in the most conservative manner possible, employing the non-parametric Kruskal-Wallis test with Dunn's correction for multiple comparisons; * $p < 0.05$, ** $p < 0.01$, *** $p < 0.001$. Bars indicate median values, boxes indicate the 25 to 75% spread, and whiskers indicate the minimum and maximum values, with individual data points shown.

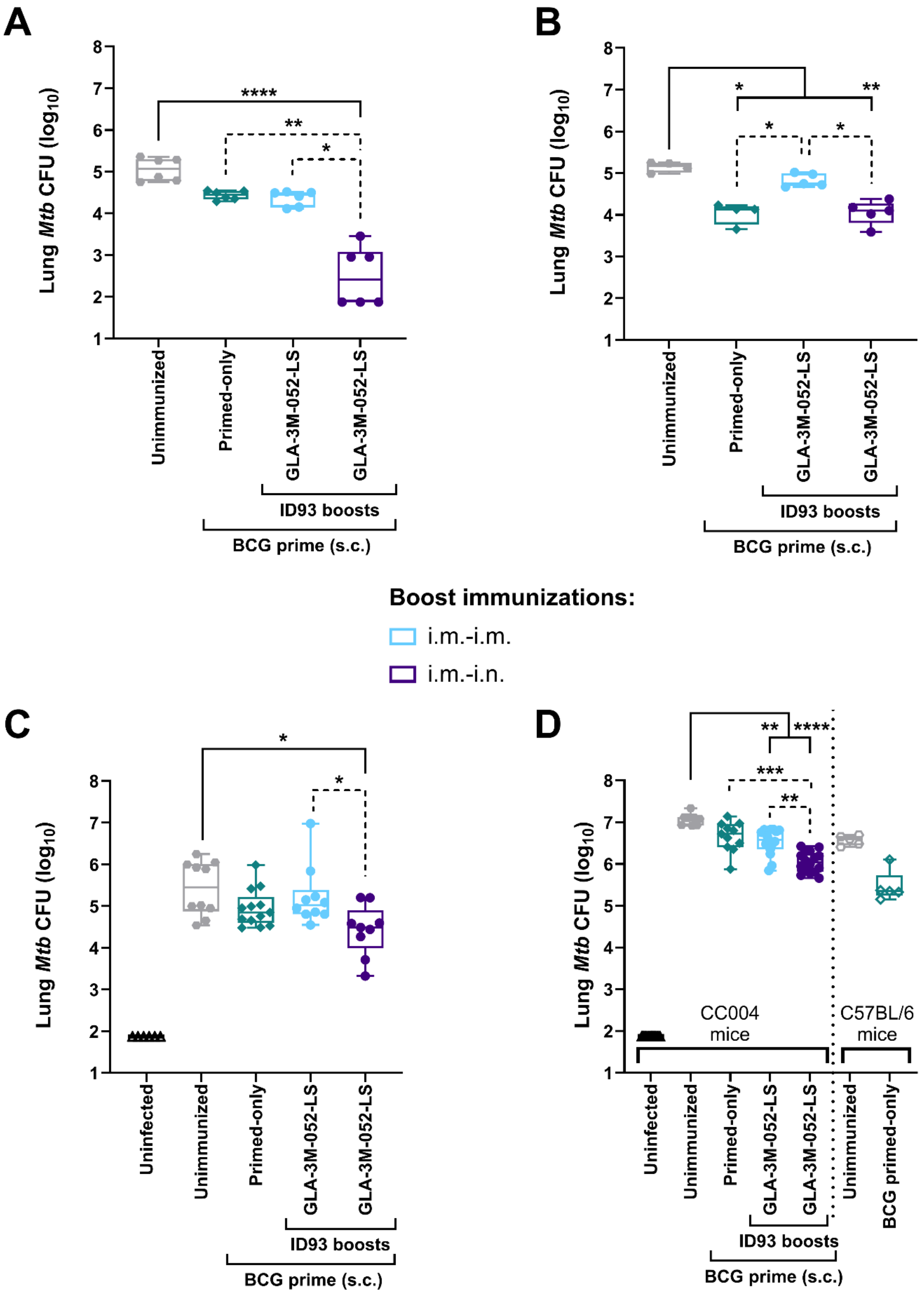
A
B
C
D
Lung Mtb CFU (log10)
Unimmunized
Primed-only
GLA-3M-052-LS
ID93 boosts
BCG prime (s.c.)
Boost immunizations:
i.m.-i.m.
i.m.-i.n.
Uninfected
CC004 mice
C57BL/6 mice
BCG primed-only

**Figure S8. Separate results from four experiments of lung CFUs in mice immunized by heterologous vaccine regimens involving BCG and ID93 + GLA-3M052-LS.** Lung CFUs ($log_{10}$) from four separate experiments involving the immunization regimens described in Table 5 and additional controls. Combined data are shown in Figure 5. Four weeks after the final immunization, mice were challenged with low-dose aerosolized *Mtb* (25 CFUs for top panels, 100 CFUs for bottom panels). Four weeks following *Mtb* challenge, lungs were harvested and CFUs measured. CFU data were log-transformed and statistical analysis was conducted using the non-parametric Kruskal-Wallis test with Dunn's correction for multiple comparisons (solid lines: BCG-primed groups compared to unimmunized group; dotted lines: comparison between all BCG-primed groups); *p<0.05, ** p<0.01, *** p<0.001, **** p<0.0001. Box-whisker plots are shown wherein bars indicate median values, boxes indicate the 25 to 75% spread, and whiskers indicate the minimum and maximum values, with individual data points shown.

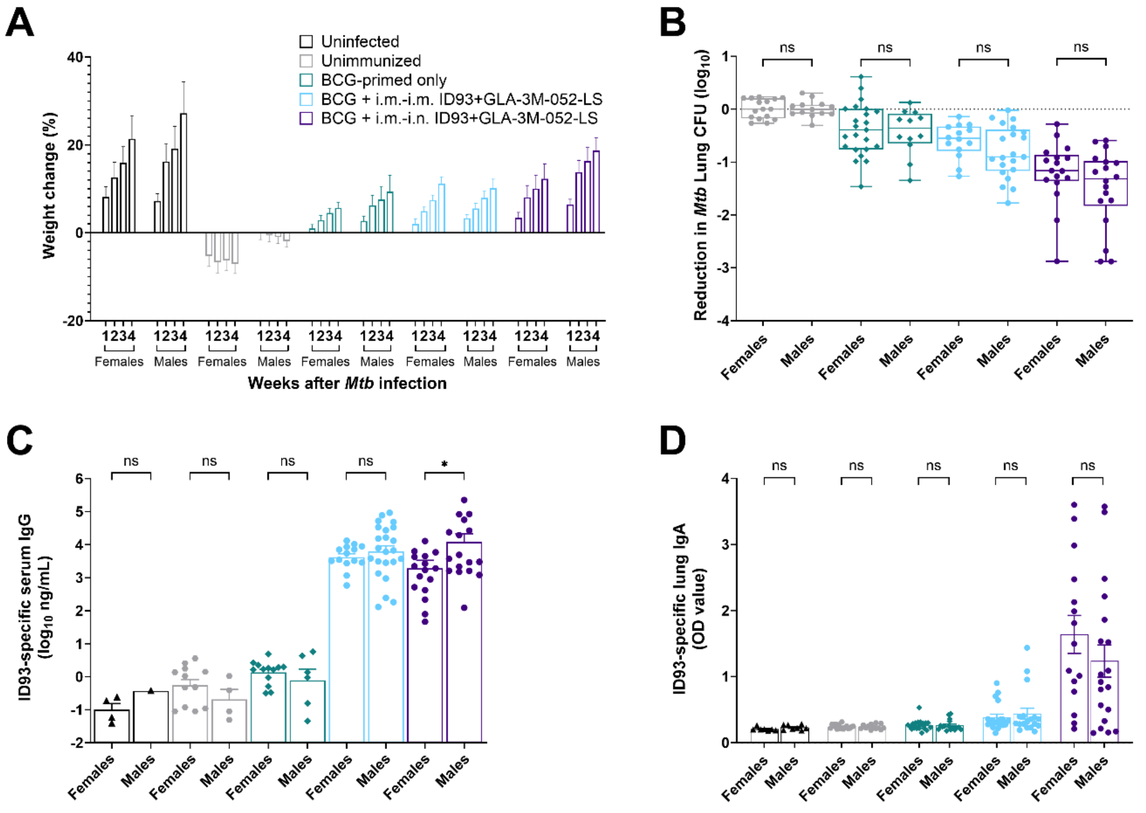


**Figure S9. Sex-specific effects in protective efficacy study using heterologous vaccine regimens in BCG-primed and ID93 + GLA-3M-052-LS boosted CC004 mice**. Combined data (total *n*=7 to 20 CC004 mice/sex/group) from the same four separate experiments shown in Supplemental Figure S8 immunized according to the regimens described in Table 5 and Figure 5. Three weeks after the final immunization, mice were challenged with low-dose (25-100 CFUs) aerosolized *Mtb*. Four weeks after *Mtb* challenge, serum and lungs were harvested and lung CFUs were quantified. **(A)** Change in body weight and **(B)** reduction in lung *Mtb* CFUs. CFU data were log-transformed and plotted as reduction in the CFUs ($\log_{10}$) compared to the average of unimmunized mice infected with *Mtb*. ID93-specific Abs from **(C)** serum IgG and **(D)** lung IgA. Bars show mean and SEM from four independent experiments. Box-and-whiskers plots show the minimum and maximum with a line at the mean. No consistent significant differences were identified between males and females within each group by unpaired *t*-test pairwise comparisons for CFU reduction, ID93-specific serum IgG, or ID93-specific lung IgA.
